# The ciliopathy protein CCDC66 controls mitotic progression and cytokinesis by promoting microtubule nucleation and organization

**DOI:** 10.1101/2022.04.22.489036

**Authors:** Umut Batman, Jovana Deretic, Elif Nur Firat-Karalar

## Abstract

Precise spatiotemporal control of microtubule nucleation and organization is critical for faithful segregation of cytoplasmic and genetic material during cell division and signaling via the primary cilium in quiescent cells. Microtubule-associated proteins (MAPs) govern assembly, maintenance, and remodeling of diverse microtubule arrays. While a set of conserved MAPs are only active during cell division, an emerging group of MAPs acts as dual regulators in dividing and non-dividing cells. Here, we elucidated the nonciliary functions and molecular mechanism of action of the ciliopathy-linked protein CCDC66, which we previously characterized as a regulator of ciliogenesis in quiescent cells. We showed that CCDC66 dynamically localizes to the spindle poles, the bipolar spindle, the spindle midzone, the central spindle and the midbody in dividing cells and interacts with the core machinery of centrosome maturation and MAPs involved in cell division. Loss-of-function experiments revealed its functions during mitotic progression and cytokinesis. Specifically, CCDC66 depletion resulted in defective spindle assembly and positioning, kinetochore fiber stability, chromosome alignment in metaphase as well as central spindle and midbody assembly and organization in anaphase and cytokinesis. Notably, CCDC66 regulates mitotic microtubule nucleation via noncentrosomal and centrosomal pathways via recruitment of gamma-tubulin to the spindle poles and the spindle. Additionally, CCDC66 bundles microtubules *in vitro* and in cells by its C-terminal microtubule-binding domain. Phenotypic rescue experiments showed that the microtubule and centrosome-associated pools of CCDC66 individually or cooperatively mediate its mitotic and cytokinetic functions. Collectively, our findings identify CCDC66 as a multifaceted regulator of the nucleation and organization of the diverse mitotic and cytokinetic microtubule arrays and provides new insight into nonciliary defects that underlie ciliopathies.

## Introduction

Faithful segregation of genetic and cytoplasmic material during cell division is essential for growth and development of multicellular organisms. Deregulation of the molecular processes that regulate cell division leads to aneuploidy and chromosomal instability and thereby to the initiation and progression of various human cancers (Lens and Medema, 2019; Tanaka and Hirota, 2009). As such, mitosis and cytokinesis are highly regulated, multistep processes involving dynamic regulation and coordinated activity of multiple cellular structures and signaling pathways (D’Avino et al., 2015; McIntosh, 2016; Verma et al., 2019). In particular, microtubule (MT) cytoskeleton undergoes a series of morphological changes to form diverse MT arrays such as the bipolar spindle, central spindle and midbody. Precise spatiotemporal control of the assembly, maintenance and dynamic remodeling of these MT arrays requires a diverse group of the MT-associated proteins (MAPs), which bind to MTs and regulate their dynamic properties, organization and stability as well as their interactions with other proteins and cellular structures (Maiato et al., 2004; Petry, 2016). Thereby, MAPs play essential roles during numerous cell cycle processes including MT nucleation, formation and organization of the mitotic spindle and central spindle, chromosome capture, alignment and segregation, cleavage furrow formation and abscission (Glotzer, 2009; Petry, 2016; Petry and Vale, 2015). Proteomic profiling of MT-based structures of dividing cells, functional screens and loss-of-function studies have identified hundreds of MAPs as regulators of mitosis and cytokinesis (Amin et al., 2019; Petry, 2016). However, key questions remain about their functions, mechanisms and links to disease as well as how they cooperate with different cellular structures (i.e. centrosomes), protein complexes and signaling pathways to modulate the parameters that ultimately define the size, shape and dynamics of MT arrays.

In animal somatic cells, MT nucleation is initiated at the centrosomes, the preexisting spindle MTs and the chromatin, with centrosomes being the major MT-organizing centers (Duncan and Wakefield, 2011; Petry and Vale, 2015). The mechanisms by which these distinct pathways work, their relative contributions to formation of diverse MT arrays in cells and the extent of their crosstalk have been an area of active investigation. As cells enter mitosis, pericentriolar material (PCM) around centrioles expands in a process called centrosome maturation, which increases its MT-nucleation capacity (Luders and Stearns, 2007; Palazzo et al., 2000). Centrosome maturation is initiated by PLK1-dependent phosphorylation of Pericentrin and CDK5RAP2, which promotes recruitment of additional PCM proteins including Cep152, Cep192 and gamma-tubulin (Cabral et al., 2019; Choi et al., 2010; Conduit et al., 2014; Dobbelaere et al., 2008; Gomez-Ferreria et al., 2007; Hanafusa et al., 2015; Joukov and De Nicolo, 2018; Lee and Rhee, 2011). Acentrosomal MT nucleation during mitosis is triggered at the chromosomes in RanGTP, Op8/Stathmin and the chromosomal passenger complex (CPC)-dependent pathways, and at the spindle MTs in a HAUS/augmin complex-dependent way (Goshima et al., 2008; Goshima et al., 2007; Heald et al., 1996; Lawo et al., 2009; Petry and Vale, 2015).

Organization of MTs into highly ordered mitotic and cytokinetic arrays play critical roles for cell division. For example, 20-40 kinetochore MTs in human cells form parallel bundles termed K-fibers, which run from spindle poles to kinetochores and are essential for chromosome alignment and segregation (Booth et al., 2011; McDonald et al., 1992; McEwen et al., 1997; Tolic, 2018). Bridging fibers, composed of antiparallel bundles of interpolar MTs, connect two sister K-fibers and push them apart to separate spindle poles (Tolic, 2018; Vukusic et al., 2017). Similarly, in anaphase, an antiparallel MT bundle forms the central spindle/midzone which pushes the spindle poles to opposite side of the cell and directs the localization of ingression furrow important for the division of cytoplasm (Glotzer, 2009). Anti-parallel MT bundles at the spindle midzone are crosslinked by the evolutionarily conserved Protein Translocator of Cytokinesis 1 (PRC1)/Ase1/MAP65 family (Bieling et al., 2010; Janson et al., 2007; Mollinari et al., 2002; Subramanian et al., 2010). Central spindle shortens to form the midbody during cytokinesis, which will direct the membrane abscission site (Hu et al., 2012). Moreover, astral MTs, which emanate from the centrosomes, interact with the cell cortex to position the spindle within a cell and determine the initial cleavage plane through communication with the equatorial cortex (D’Avino et al., 2005). Although multiple MAPs involved in the formation and stabilization of the distinct spindle bundles have been identified, the full extent of MAPs involved in MT crosslinking and stability as well as their mechanism of action have yet to be determined in future studies.

The drastic remodeling of the MT network during cell division requires precise regulation of when and where MAPs are activated. A subset of MAPs is active during mitosis but inactive during interphase. Such regulation is achieved by modulation of their affinity to MTs, regulation of their cellular abundance and localization and posttranslational modifications (Syred et al., 2013). Importantly, there is also a group of MAPs with dual functions in dividing and non-dividing cells (Akhmanova and Steinmetz, 2015; Kaplan et al., 2001; Villari et al., 2020). For example, End Binding 1 (EB1) regulates MT plus-end dynamic and targets other MAPs to the plus ends both in interphase and mitosis (Green et al., 2005; Schroder et al., 2011; Schroder et al., 2007; Song et al., 2021). Recently, a critical regulator of primary cilium assembly in non-dividing cells, Intraflagellar transport Protein 88 (IFT88), has been described for its functions during mitotic spindle orientation and central spindle organization (Delaval et al., 2011; Taulet et al., 2019). Importantly, the discovery of non-ciliary functions of IFT88 unraveled that its mutations might contribute to polycystic kidneys with both impaired ciliary function and aberrant cell division (Delaval et al., 2011; Taulet et al., 2019). Despite the progress made in the characterization of MAPs with dual roles in cycling and non-cycling cells, questions remain about their functions, mechanisms, and modes of regulation in different stages of the cell cycle.

We previously characterized coiled coil protein 66 (CCDC66) as a MAP and a regulator of primary cilium formation and composition in quiescent cells(Conkar et al., 2019; Conkar et al., 2017). It was originally described as a gene mutated in retinal degeneration and later characterized for its retinal and olfactory functions using CCDC66^-/-^ mouse (Dekomien et al., 2010; Gerding et al., 2011; Murgiano et al., 2020; Schreiber et al., 2018). Recently, CCDC66 was identified as part of the Joubert syndrome interaction network consisting of other MAPs such as CSPP1, TOGARAM1 and CEP290 (Latour et al., 2020). Consistent with its link to ciliopathies, we and others previously showed that retinal degeneration mutations disrupt its ciliary functions and interactions (Conkar et al., 2017; Murgiano et al., 2020). In addition to its ciliary functions, following lines of evidence suggest that CCDC66 might function as a regulator of cell division: CCDC66 mRNA was identified in the MT-interacting transcriptome of *Xenopus tropicalis*, indicative of its functions during MT-based cellular processes (Sharp et al., 2011). Additionally, CCDC66 localized to spindle poles and MTs in dividing cells and its depletion led to disorganized poles in mitotic cells (Conkar et al., 2017; Sharp et al., 2011). Finally, CCDC66 proximity interactome generated from asynchronous cells revealed interactions with regulators of cell division (Conkar et al., 2017; Gheiratmand et al., 2019; Gupta et al., 2015). However, the full extent of CCDC66 functions during different stages of cell division and the underlying molecular mechanisms are not known. Addressing these key unknowns will uncover the relationship of CCDC66 with other components of the mitotic and cytokinetic machinery and also provide insight into whether and if so, how its nonciliary functions contribute to its disease mechanisms.

In this study, we examined the localization, interactions, functions, and mechanisms of CCDC66 during cell division.

## Results

### CCDC66 localizes to spindle poles and MTs during mitosis and cytokinesis

Based on its previously reported localization and interactions, we hypothesize that CCDC66 plays important roles during mitosis and cytokinesis (Conkar et al., 2017; Gheiratmand et al., 2019; Sharp et al., 2011). To test this, we first examined localization of endogenous and mNG-CCDC66 fusion proteins at different cell cycle stages in mammalian cell lines. Antibody against endogenous CCDC66 revealed its localization to the centrosome throughout the cell cycle (Fig. 1A, S1A). In dividing cells, CCDC66 also localized to multiple MT-based structures including the spindle MTs in prometaphase and metaphase, the central spindle in anaphase and the midbody in cytokinesis in human osteosarcoma (U2OS) cells (Fig. 1A, S1A). To examine the dynamic localization of CCDC66 during cell cycle, we generated cell lines that stably express mNeonGreen (mNG)-CCDC66 using lentiviral transduction. mNG protein was chosen over GFP as the fluorescent tag due to its higher fluorescent intensity and stability (Shaner et al., 2013). Stable expression of the fusion protein in U2OS and RPE1 cells was validated by immunoblotting using mNG antibody and immunofluorescence using anti-CCDC66 antibody (Fig. S1B-D). In fixed U2OS and RPE1 stable cells, mNG-CCDC66 localized to the centrosome and centriolar satellites in interphase cells (Fig. 1B, S1E) and to the spindle poles and MT-based structures of mitosis and cytokinesis in dividing cells (Fig. 1B, S1E). Although CCDC66 localization to the astral MTs and midzone was apparent in cells stably expressing mNG-CCDC66, it was very weak in cells stained for endogenous CCDC66 (Fig. 1A and S1A). This might be due to the high cytoplasmic and punctate background associated with CCDC66 antibody staining and/or the relatively lower abundance of CCDC66 at the astral microtubules and spindle midzone. In agreement with its localization in fixed cells, time-lapse imaging of mNG-CCDC66 cells stained with SIR-tubulin showed that CCDC66 dynamically localized to the centrosome, centriolar satellites and MT-based structures in different cell states (Fig. S1F, S1G, Movie 1, Movie S1).

**Figure 1.**
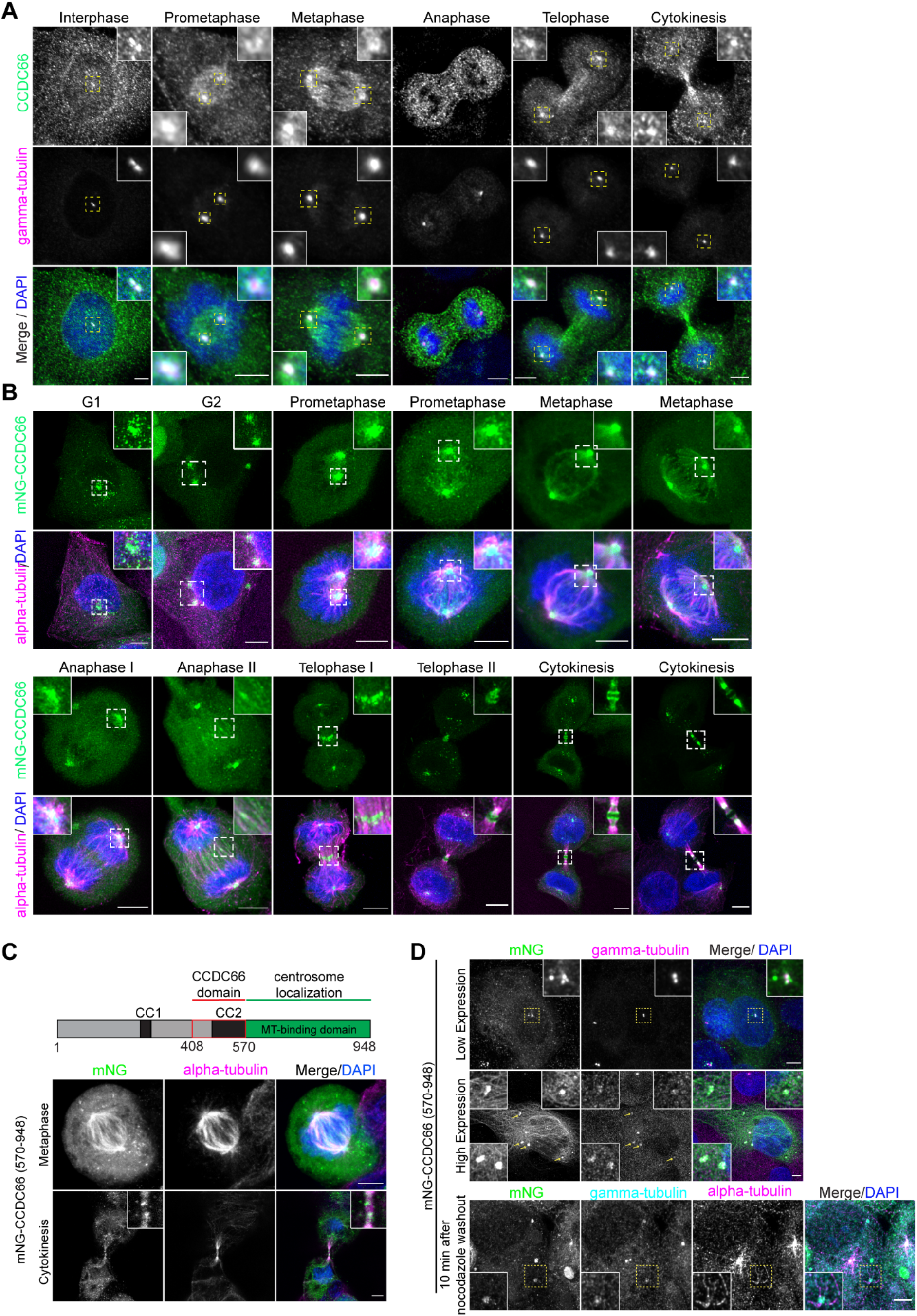
CCDC66 localizes to spindle poles and microtubule-based structures during cell division. (A) Localization of CCDC66 at different stages of the cell cycle. U2OS were fixed with methanol followed by acetone and stained for CCDC66, gamma-tubulin and DAPI. Scale bar: 5 µm, insets show 4X magnifications of the boxed regions. (B) Localization of mNeonGreen-CCDC66 at different stages of the cell cycle. U2OS cells stably expressing mNeonGreen-CCDC66 fusion (U2OS::mNG-CCDC66) were fixed with 4% PFA and stained for alpha tubulin and DAPI. Scale bar: 5 µm, insets show 4X magnifications of the boxed regions. (C) Schematic representation of full length (FL) CCDC66 domain organization. CC1 and CC2 indicate coiled-coil domains. The C-terminal region was previously described as a microtubule-binding region (Conkar et al., 2017). Still confocal images show U2OS cells transfected with mNeonGreen-CCDC66 C-terminal construct (mNG-CCDC66^570-948^). 24 h post-transfection, cells were fixed with 4% PFA and stained for alpha-tubulin and DAPI. mNG-CCDC66^570-948^ localizes to spindle poles and spindle microtubules during metaphase and to midbody during cytokinesis. Scale bar: 5 µm. (D) mNG-CCDC66^570-948^ sequesters gamma-tubulin to cytoplasmic aggregates. U2OS cells were transfected with mNG-CCDC66^570-948^, fixed with 4% PFA and stained for gamma-tubulin and DAPI. Top panel shows a lower expressing cell in which mNG-CCDC66^570-948^ is restricted mostly to centrosomes. Middle panel shows a high-expressing cell with multiple cytoplasmic aggregates of mNG-CCDC66^570-948^ that co-localize with gamma-tubulin. Bottom panel shows mNG-CCDC66^570-948^ transfected cells that are treated with 5 µg/ml nocodazole for 1 h at 37 °C and incubated for 10 min after nocodazole washout. Scale bar: 5 µm, insets show 4X magnifications of the boxed regions.

We showed that CCDC66 is required for recruitment of core machinery of centrosome maturation to the spindle poles and acts as a bundling protein *in vitro* and in cells. Its association with centrosomes and MTs is required for spindle assembly and organization, k-fiber and midbody integrity, chromosome alignment, and cytokinesis. Our findings unravel non-ciliary functions for CCDC66 during cell division and provides insight into the integrated activity of centrosomes and MAPs during spatiotemporal regulation of MT nucleation and organization in mitosis and cytokinesis.

Previously, we showed that CCDC66 and its C-terminal 570-948 residues bind to MTs *in vitro* (Conkar et al., 2017). Given its direct microtubule affinity, we tested whether this C-terminal fragment recapitulates the localization of full length CCDC66 during cell division. Like mNG-CCDC66, mNG-CCDC66 (570-948) localized to the spindle poles, bipolar spindle and central spindle during mitosis and midbody during cytokinesis in U2OS cells (Fig. 1C). When overexpressed, mNG-CCDC66 (570-948) formed cytoplasmic aggregates that **CCDC66 co-localizes and interacts with critical regulators of mitosis and cytokinesis** recruited gamma-tubulin, suggesting a putative interaction between them (Fig. 1D). To investigate the functional significance of this recruitment during MT nucleation, we performed MT regrowth experiments and found that the cytoplasmic aggregates nucleated MTs 10 min after nocodazole washout (Fig. 1D). Together, these data indicate that the C-terminal 379 residues of CCDC66 is sufficient for its cellular localization to the centrosome and MTs in dividing cells.

We defined the high-resolution localization of CCDC66 relative to the markers of the spindle poles, bipolar spindle, central spindle and midbody. Specifically, we co-stained U2OS::mNG-CCDC66 cells with antibodies against PCM proteins gamma-tubulin, CEP192, CEP152, CDK5RAP2 (spindle poles), the MT-crosslinking protein PRC1 (central spindle and midbody MTs), the centrosome protein CEP55 (midbody core), the kinesin motor protein Kif23/MKLP1and phosphorylated Aurora A/B/C (midbody flanks) (Fry et al., 2017; Glotzer, 2009; Hu et al., 2012). CCDC66 localized to the spindle poles throughout mitosis, as shown by its co-localization with gamma-tubulin, CDK5RAP2, CEP192 and CEP152 (Fig. 2A, 2B). Notably, the spindle pole-associated pool of CCDC66 was maintained in nocodazole-treated cells, confirming that this pool binds to the spindle poles independent of MTs (Fig. 2C). To determine the precise cytokinetic localization of CCDC66, we performed plot profile analysis along the midbody MT bundles for CCDC66 and known midbody markers (Fig. 2D) (Hu et al., 2012). In cytokinesis, CCDC66 localized to two closely spaced bands at the midbody. Plot profile analysis revealed its co-localization with PRC1 at the dark zone (Fig. 2D), which is defined as the narrow region on the MT bundle in the center of the midbody (Hu et al., 2012). mNG-CCDC66 also co-localized with PRC1 at the spindle midzone in anaphase and telophase cells (Fig. S2A). In contrast, CCDC66 did not co-localize with CEP55 and MKLP1 at the midbody, which are visualized as rings representing the bulge at the center of the midbody, and phospho-Aurora (Fig. 2D), which marks the broader bands on MTs outside the dark zone (Hu et al., 2012). Of note, we observed accumulation of CCDC66 outside the midbody in cells at later stages of cytokinesis, which is also evident in the dynamic behavior of mNG-CCDC66 during cell division (Fig. 2D, Movie 1, Movie S1). These CCDC66-positive structures did not co-localize with the known midbody markers and were not detected in cells stained for endogenous CCDC66 (Fig. 1A).

**Figure 2.**
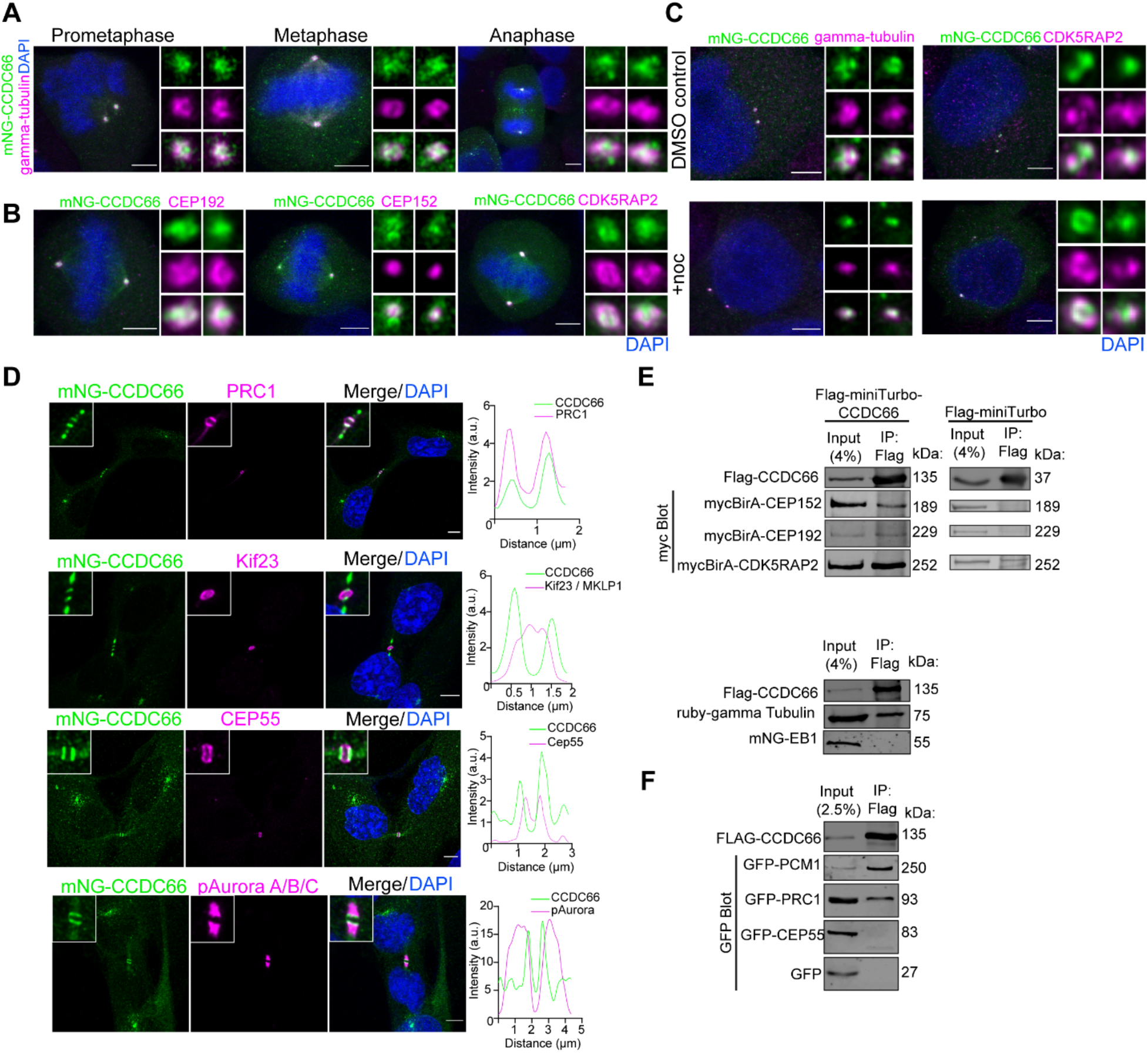
CCDC66 co-localizes and interacts with PCM proteins and spindle/midbody MAPs. (A) mNG-CCDC66 localizes to the centrosome throughout the cell cycle. U2OS::mNG-CCDC66 cells were fixed with 4% PFA and stained for gamma-tubulin and DAPI. Still deconvolved confocal images represent centrosomal co-localization of mNG-CCDC66 with gamma-tubulin during different stages of cell division. Scale bar: 5 µm, insets show 4X magnifications of the centrosomes. (B) Localization of mNG-CCDC66 in U2OS cells relative to Pericentriolar Material (PCM) proteins. U2OS::mNG-CCDC66 cells were fixed with 4% PFA and stained for CEP192, CEP152, or CDK5RAP2 and DAPI. Deconvolved confocal images represent their relative localization at the spindle poles during metaphase. Scale bar: 5 µm, insets show 4X magnifications of the centrosomes. (C) Effect of microtubule depolymerization on CCDC66 localization at the centrosome. U2OS::mNG-CCDC66 cells were treated with 0.1% DMSO or 5 µg/ml nocodazole for 1h. Cells were then fixed with 4% PFA and stained for gamma-tubulin or CDK5RAP2 and DAPI. Scale bar: 5 µm, insets show 4X magnifications of the centrosomes. (D) Localization of mNG-CCDC66 in RPE1 cells relative to midbody proteins. RPE1::mNG-CCDC66 cells were fixed with methanol and stained for mNeonGreen and PRC1, Cep55, Kif23 (MKLP1) or pAurora A/B/C and DAPI. Graphs show the plot profiles to assess co-localization with the indicated marker. Using ImageJ, a straight line was drawn on the midbody and intensity along the distance was plotted on Graphpad Prism. (E) Co-immunoprecipitation of Flag-miniTurbo-CCDC66 or Flag-miniTurbo with PCM proteins from HEK293T cells. Flag-minTurbo-CCDC66 was co-transfected with myc-BirA-Cep192, myc-BirA-Cep152 and myc-BirA-CDK5RAP2 in HEK293T cells. Flag-miniTurbo-CCDC66 and Flag-miniTurbo was precipitated with Flag beads and input and eluates (IP) were blotted with Flag and myc antibodies to assess the interaction. HEK293T cells were also co-transfected with Flag-miniTurbo-CCDC66 and Ruby-Gamma-tubulin-T2A-mNG-EB1 fusion construct. Flag-miniTurbo-CCDC66 was precipitated with Flag beads and Input and eluates (IP) were blotted with Flag, EB1 and gamma tubulin antibodies. (F) Co-immunoprecipitation of Flag-CCDC66 with midbody proteins from HEK293T cells. Flag-CCDC66 was co-transfected with GFP, GFP-Cep55 or GFP-PRC1 in HEK293T cells. Flag-CCDC66 was precipitated with Flag beads and Input and eluates (IP) were blotted with Flag and GFP antibodies to assess the interaction.

To generate hypothesis regarding the functions and mechanisms of CCDC66 during cell division, we compiled a list of CCDC66 interactors from high throughput proximity-mapping studies and performed Gene Ontology (GO)-enrichment analysis of its interactors. (Fig. S2B, S2C) (Conkar et al., 2017; Gheiratmand et al., 2019; Gupta et al., 2015). In addition to proteins involved in primary cilium assembly and function, CCDC66 proximity interactors were enriched for biological processes related to cell division including spindle assembly and organization, chromosome segregation, metaphase plate congression and cytokinesis (Fig. S2B). Consistently, cellular compartments involved in these processes such as the HAUS complex, mitotic spindle, spindle pole and kinetochore were among the enriched GO categories (Fig. S2C). This analysis supports nonciliary functions for CCDC66 during cell division.

To define the physical mitotic and cytokinetic interactions of CCDC66, we performed Flag-based co-immunoprecipitation experiments with PCM proteins and mitotic MAPs that co-localize with CCDC66 (Fig. 2A-D) or were described as critical regulators of mitosis and cytokinesis. As a positive control for interaction experiments, we used the centriolar satellite marker protein PCM1, which we previously reported as an interactor for CCDC66 (Conkar et al., 2017). First, we assayed whether CCDC66 interacts with critical regulators of centrosome maturation. CCDC66 co-pelleted with myc-BirA* fusions of CDK5RAP2, Cep192, Cep152 and gamma-tubulin (Fig. 2E). As negative controls, FLAG-miniTurbo did not co-pellet with myc-BirA* fusions of these positive interactions, and CCDC66 did not co-pellet with the MT plus-end-tracking protein EB1 (Fig. 2E). Next, we tested interactions of CCDC66 with regulators of cytokinesis (Glotzer, 2009). Flag-miniTurbo-CCDC66 interacted with the GFP-PRC1, but not GFP-CEP55 and the negative control GFP (Fig. 2F). Together with its localization profile during cell division, the new CCDC66 interactors we identified suggest potential mitotic and cytokinetic functions of CCDC66 via regulating centrosome maturation, MT nucleation and/or organization.

### CCDC66 is required for mitotic progression and cytokinesis

To elucidate CCDC66 functions during mitotic and cytokinetic progression, we performed loss-of-function experiments using an siRNA validated in depletion and rescue experiments (Conkar et al., 2017). Given that CCDC66 localization and dynamics in dividing cells were similar in both RPE1 and U2OS cells (Fig. 1, S1), we chose U2OS cells for further characterization as a p53-responsive transformed cell (McKinley and Cheeseman, 2017). Immunoblotting and immunofluorescence analysis of U2OS cells 48 h after transfection with control and CCDC66 siRNAs confirmed efficient depletion (Fig. S3A, S3B). To investigate the role of CCDC66 in the dynamic events of the cell cycle, we performed time lapse confocal imaging of control and CCDC66-depleted U2OS cells that stably express the chromosome marker mCherry-H2B and determined the fate of the dividing cells (Fig. 3A). We plotted the fate of individual control and CCDC66-depleted cells as vertical bars in Fig. S2C, where the height of the bar represents the mitotic time and the color of the bars represent the different fates (gray: successful division, pink: mitotic arrest, cyan: apoptosis). The mitotic time, which was defined as the time from nuclear envelope breakdown to anaphase, increased about 1.4-fold in CCDC66-depleted cells relative to control cells (siCCDC66: 39.2 ± 8.6 min, siControl: 27.1 ± 6.6 min) (Fig. 3B, Movies 2, Movie 3). The mitotic cells that did not complete mitosis during 12 h of live imaging had two different fates. They either exhibited prometaphase arrest, with some cells reaching chromosome alignment followed by metaphase plate regression or underwent apoptosis, which was assessed by membrane blebbing and DNA fragmentation (Fig. 3A, 3C, 3D, Movies 4-6). As compared to 8.8 ± 3.6% of control cells, 23.3 ± 3.7% percent of CCDC66-depleted cells died after prolonged mitosis (Fig. 3D). We also quantified the mitotic index by scoring the percentage of cells positive for the mitotic marker phospho-H3 and found that it was not altered upon CCDC66 loss (Fig. S3D). This might be due to increased apoptosis in U2OS cells as a response to the delayed mitosis followed by mitotic failure.

**Figure 3.**
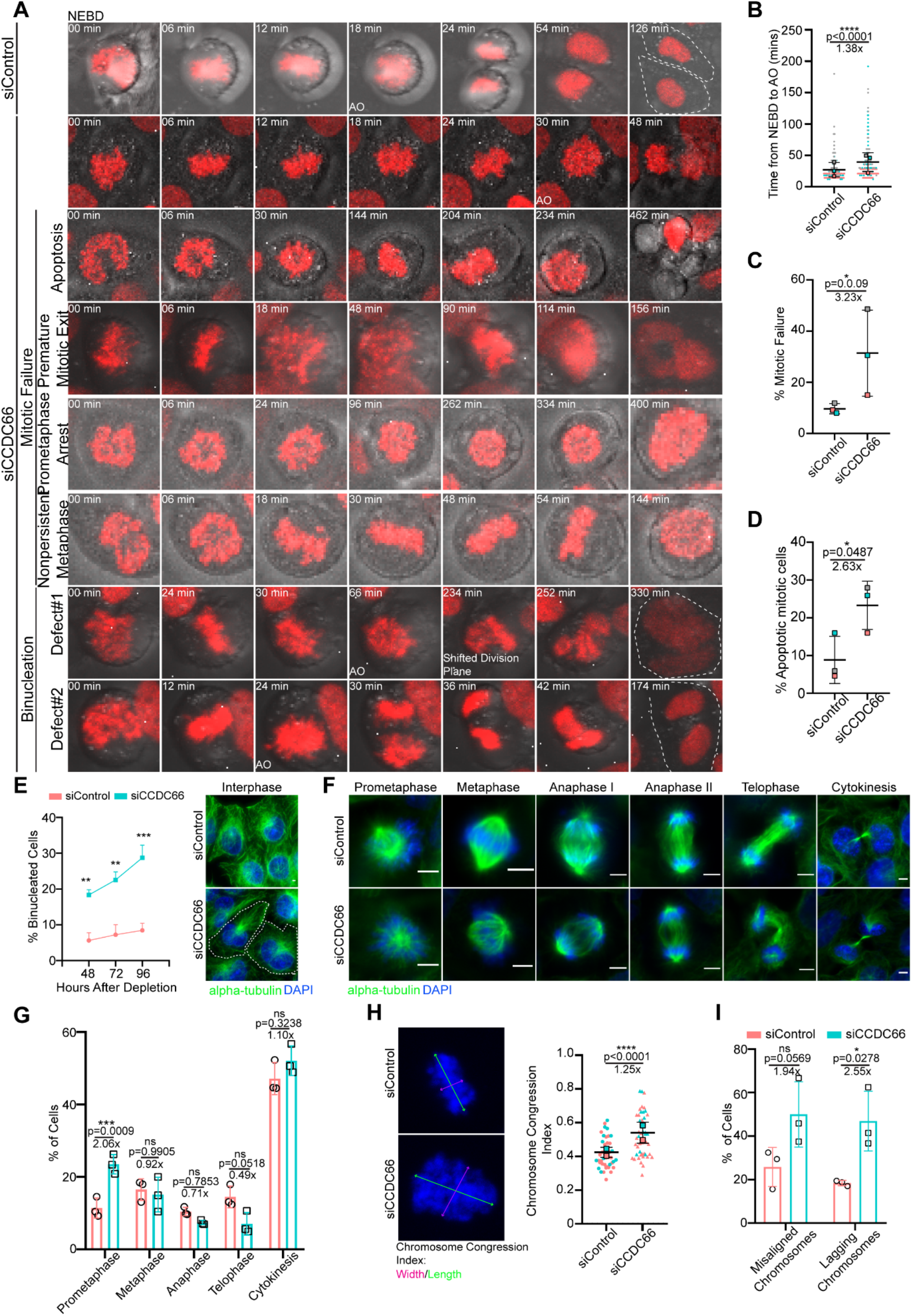
CCDC66 is required for faithful mitotic progression and cytokinesis. (A) Effects of CCDC66 depletion on mitotic progression. U2OS::mcherry-H2B cells were grown on 2-well Lab-Tek plates and transfected with non-targeting (siControl) or CCDC66 targeting (siCCDC66) siRNA. After 48h of transfection, cells were imaged with confocal microscopy with 20x objective. Images are acquired every 6 mins for 12h. Representative still images from live imaging are shown for different phenotypic categories for siCCDC66 include apoptosis, premature mitotic exit, prometaphase arrest and non-persistent metaphase. (B) Quantification of mitotic time from A. Mitotic time was quantified as the time interval from nuclear envelope breakdown (NEBD) to anaphase onset (AO). Data represents the mean ±SEM of three independent experiments and is plotted using super-plot. (C) Quantification of percent mitotic failure from A. Mitotic failure refers to the cells that entered mitosis but could not proceed through anaphase because of; premature mitotic exit, prometaphase arrest and non-persistent metaphase. Data represents the mean ±SEM of three independent experiments and is plotted using super-plot. (D) Quantification of percent mitotic cell death from A. Mitotic cell death represents mitotic cells that underwent apoptosis during the time of imaging. Data represents the mean ±SEM of three independent experiments and is plotted using super-plot. (E) CCDC66 depletion increases binucleation. Cells were transfected with either siControl or siCCDC66, fixed with methanol 48, 72, and 96h post-transfection and stained for alpha-tubulin and DAPI. Binucleation was categorized based on DNA and alpha tubulin staining. Scale bar: 5 µm (F) Spindle and chromosome phenotypes of CCDC66-depleted cells. Representative images of different stages of cell division for siControl and siCCDC66 transfected U2OS cells are shown. Cells were transfected with either siControl or siCCDC66, fixed with methanol 48h post-transfection and stained for alpha-tubulin and DAPI. Mitotic cell stages were categorized based on DNA staining. Scale bar: 5 µm (G) Quantification of E. Data represents mean ±SEM of three independent experiments. n>1000 for all experiments. Mean prometaphase percentage is 11.42 for siControl and mean prometaphase percentage is 23.50 for siCCDC66. (***p<0.001, ns not significant) (H) CCDC66 depletion increases chromosome width. U2OS cells were transfected with siRNA, fixed with methanol and stained for DAPI. Chromosome congression index is measured by dividing the length of the metaphase plate by its width. Data represent the mean ±SEM of two independent experiments. (****p<0.0001). Scale bar: 5 µm. (I) Quantification of chromosome alignment and segregation defects from A. The quantification shows the number of metaphase cells that have misaligned chromosomes and anaphase and telophase cells that have lagging chromosomes. Data represent the mean ±SEM of three independent experiments. (ns not significant).

In addition to mitotic defects, we observed that CCDC66 depletion interfered with progression of cytokinesis. Time lapse imaging of dividing CCDC66-depleted cells revealed two different events that resulted in binucleated cells (Fig. 3A, Movies 7,8). First phenotype was the regression of the cleavage furrow after chromosome segregation. Second phenotype was failure in forming the cleavage furrow and initiating cytokinesis (Fig. 3A, Movie 7, Movie 8). In a complementary approach, we quantified the percentage of binucleated cells in a time course manner by fixing cells at different time points after siRNA transfection. CCDC66 depletion caused an increase in the percentage of binucleated cells in a time course manner (Fig. 3E). These results identify CCDC66 as a regulator of cytokinesis.

To further define its functions during mitotic progression, we examined whether CCDC66 depletion leads to the accumulation of cells at a particular stage of mitosis by scoring cells based on their chromosome positioning and spindle organization using DNA and MT staining (Fig. 3F) (Baudoin and Cimini, 2018). CCDC66 depletion led to a higher fraction of cells in prometaphase, suggesting the presence of chromosome and/or spindle-related defects (Fig. 3F, 3G). In agreement, we noted that the MT arrays of the bipolar spindle, central spindle and midbody of CCDC66-depleted cells were disorganized (Fig. 3F). To quantify chromosome alignment defects, we measured the ratio of the width to the height of the chromosomal mass, which was previously described as the chromosome congression index (Fig. 3H) (Green and Kaplan, 2003; Huhn et al., 2017). CCDC66-depleted cells had a higher chromosome congression index than control cells, which indicates defective chromosome alignment at the metaphase plate (Fig. 3H). Moreover, the percentage of cells with lagging chromosomes, but not with misaligned chromosomes, increased in CCDC66-depleted cells relative to control cells (Fig. 3I). Taken together, our findings demonstrate that CCDC66 regulates mitotic progression and cytokinesis in part via ensuring proper chromosome alignment and spindle assembly.

### CCDC66 is required for spindle assembly and positioning

The localization of CCDC66 to spindle microtubules suggest that mitotic progression defects associated with its depletion might be a consequence of defective spindle assembly and positioning. To investigate this, we analyzed various spindle properties in control and CCDC66-depleted cells. First, we measured the angle between the spindle axis and the substratum, which revealed an increase from 7.7 ± 0.3º in control cells to 13.1 ± 1.5º in CCDC66-depleted cells (Fig. 4A). By taking into account the changes into spindle angle, we quantified spindle length and found that it was not altered upon CCDC66 depletion (Fig. 4A). Likewise, spindle pole width was comparable between control and CCDC66-depleted cells (Fig. 4A). The essential role of astral MTs in spindle positioning led us to investigate CCDC66 functions during astral MT assembly and stability (Lu and Johnston, 2013). To this end, we quantified the astral MT fluorescence intensity and length in control and CCDC66-depleted cells and found that they were both reduced upon CCDC66 loss (Fig. 4B, 4C).

**Figure 4.**
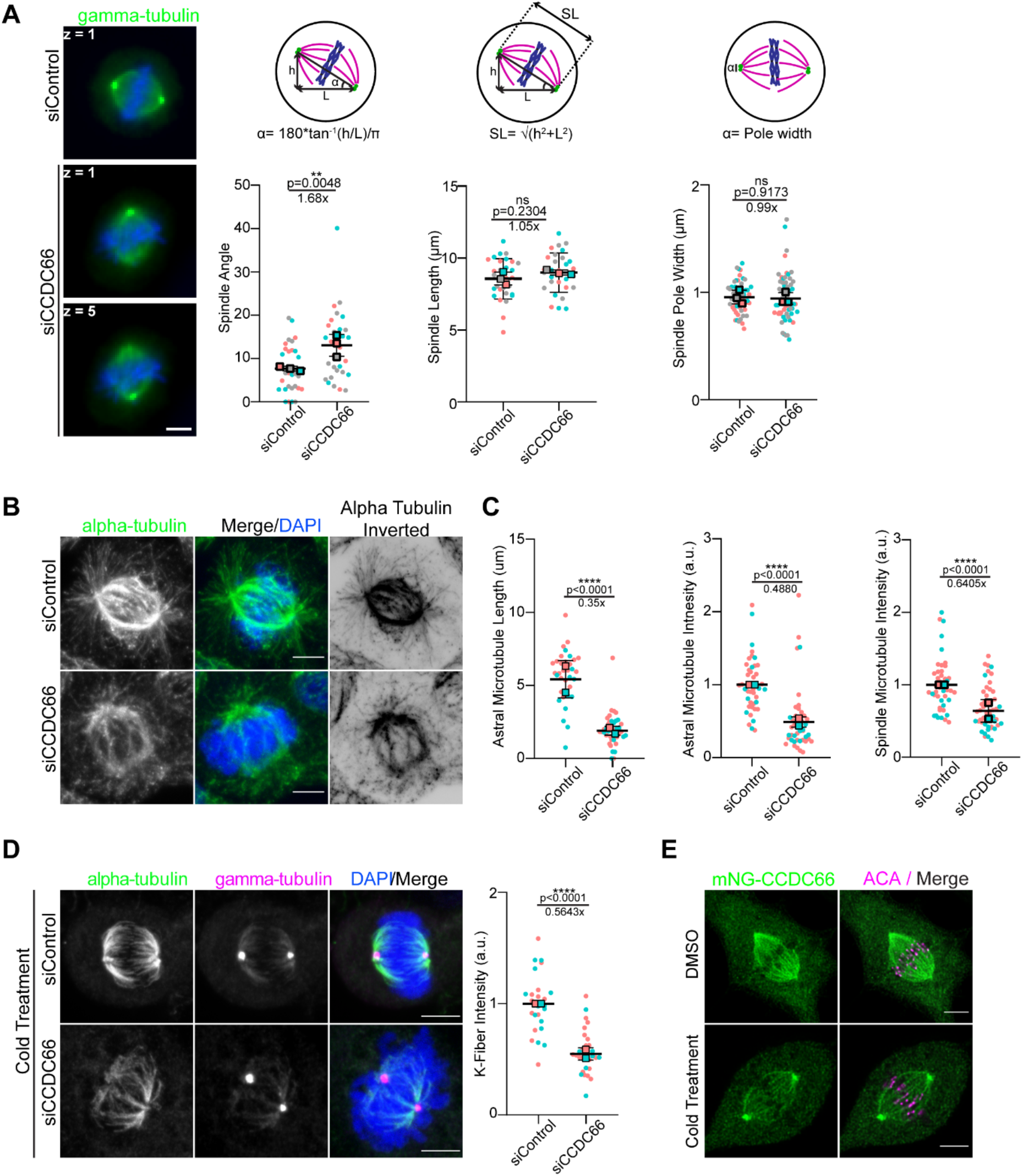
CCDC66 regulates spindle organization and positioning. (A) Effects of CCDC66 depletion on spindle angle, length and pole width. U2OS cells were transfected with control and CCDC66 siRNA, fixed with methanol and stained for gamma-tubulin. z on still images indicates the stack where the spindle pole is found. Spindle angle is calculated by the formula α=180*tan^-1^(h/L)/π where h represents the stack difference between two spindle poles, L represents the distance between spindle poles. Spindle angle is calculated by the formula SL= √(h2+L2) where h represents the stack difference between two spindle poles, L represents the distance between spindle poles. Spindle pole width is calculated by measuring the length of the pericentriolar material of the spindle pole. Data represent the mean ±SEM of three independent experiments. (**p<0.01, ns not significant). Scale bar: 5 µm. (B) Spindle microtubule density and astral microtubule length is reduced in CCDC66 depleted cells. U2OS cells were transfected with siRNA then fixed with methanol and stained for alpha-tubulin and DAPI. Representative images are shown. Inverted image is shown to emphasize astral microtubules better. Scale bar: 5 µm. (C) Quantification of (B). Astral microtubule and spindle microtubule intensity were measured on ImageJ by taking several points on the spindle to measure the intensity and subtracting the background mean intensity. Data represent the mean ±SEM of three independent experiments. (****p<0.0001). (D) CCDC66 depletion reduces K-fiber intensity. Cells were transfected with siRNA then 48 h after transfection, cells were incubated in ice for 10 mins. Cells were fixed with methanol and stained for alpha tubulin, gamma tubulin, and DAPI. K-fiber intensity was measured as described for (C). Data represent the mean ±SEM of two independent experiments. (****p<0.0001). Scale bar: 5 µm. (E) CCDC66 localizes to K-fibers. RPE1::mNG-CCDC66 stable cell line was incubated in ice for 10 mins and fixed with MeOH then stained for mNG and anticentromeric antibody (ACA). Images represent a single stack and were captured with the same camera settings from the same coverslip. Scale bar: 5 µm.

Likewise, the tubulin fluorescence intensity at the spindle in metaphase cells decreased about 0.6-fold in CCDC66-depleted cells relative to control cells (Fig. 4B, 4C). Immunoblotting of lysates prepared from control and CCDC66 siRNA-transfected cells showed that the intensity changes in spindle MTs were not due to altered cellular abundance of alpha-tubulin (Fig. S4A). Together, these results show that CCDC66 is required for spindle MT assembly and stability.

Proper formation and organization of K-fibers is essential for chromosome alignment and segregation, suggesting that chromosome-related defects in CCDC66-depleted cells could be due to impaired K-fibers (Dumont and Mitchison, 2009; Tolic, 2018). To determine whether CCDC66 is specifically required for the K-fiber stability, we performed cold stability assay in control and CCDC66-depleted cells. In this assay, K-fibers are visualized by selective depolymerization of less stable interpolar and astral MTs by cold treatment of cells for 10 minutes at 4°C (Brinkley and Cartwright, 1975; Rieder, 1981). The tubulin fluorescence intensity of cold-stable K-fibers was reduced in CCDC66-depleted cells relative to control cells(Fig. 4D), which identify CCDC66 as a regulator of K-fiber stability. Notably, mNG-CCDC66 and endogenous CCDC66 still localized to the spindle microtubules in cold-treated cells, which confirms its localization to K-fibers (Fig. 4E, S4C). Collectively, our findings indicate that CCDC66 regulates spindle assembly and positioning to ensure proper mitotic progression.

### CCDC66 is required for the assembly and organization of the central spindle and cleavage furrow

CCDC66 localizes to the central spindle, intercellular bridge/midbody and K-fibers, which are highly organized and stable MT bundles. Moreover, it interacts and co-localizes with PRC1, a well-characterized regulator of MT bundling during cell division (Mollinari et al., 2002). These lines of data suggest that CCDC66 might regulate assembly and organization of MTs at the central spindle and cleavage furrow. To test this, we examined how loss of CCDC66 affects assembly and organization of these MT arrays in dividing cells.

Quantification of the MT fluorescence intensity along the pole-to-pole axis in anaphase cells showed that MT intensity was reduced at the central spindle in CCDC66-depleted cells (Fig. 5A, 5B). This result is analogous to the effects of CCDC66 depletion on spindle MT intensity of metaphase cells. Strikingly, CCDC66 depletion also severely disrupted the organization of central spindle MTs (Fig. 5A, 5C). Majority of control cells (80.68%) had the typical central spindle organization characterized by two dense arrays of tightly packed MTs separated by a thin line at the cell center (Fig. 5A) (Lioutas and Vernos, 2013). In contrast, a significantly higher fraction of CCDC66-depleted cells exhibited highly disorganized central spindles characterized by reduced and poorly aligned MTs (siControl: 20.32%, siCCDC66: 57.04%) (Fig. 5A, 5C).

**Figure 5.**
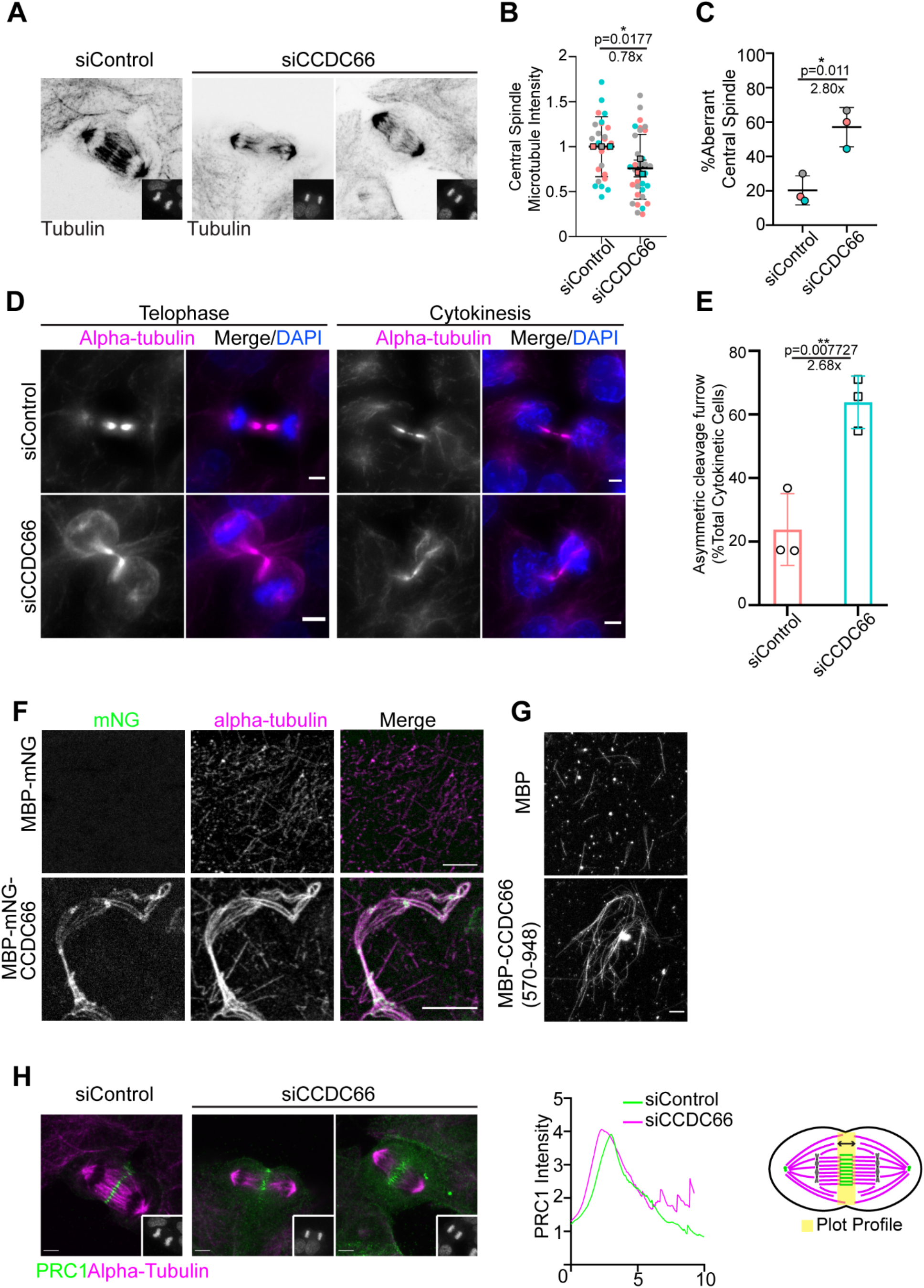
CCDC66 bundles microtubules and is required for assembly and organization of the central spindle and cleavage furrow. (A) Effect of CCDC66 depletion on central spindle assembly and organization. U2OS cells were transfected with control or CCDC66 siRNA, fixed with methanol followed 48 h post-transfection and stained for alpha tubulin and DAPI. Representative images show cells at late anaphase, as indicated by the DNA staining in the inset. (B) Quantification of (A). Graph represents the microtubule density at central spindle. Central spindle microtubule intensity was measured on ImageJ by taking several points on the spindle to measure the intensity and subtracting the background mean intensity. Data represent the mean ±SEM of three independent experiments. (*p<0.05). (C) Quantification of (A). Graph represents the percentage of aberrant central spindle with mean ±SEM of two independent experiments. (*p<0.05). Aberrant central spindle is characterized by reduced and poorly aligned microtubules. (D) CCDC66 depletion impairs the geometry of the cleavage furrow. U2OS cells were transfected with the indicated siRNAs, fixed with methanol and stained for alpha-tubulin and DAPI. Images represent telophase and cytokinetic cells. Scale bar: 5 µm. (E) Quantification of (D). Asymmetric cleavage furrow indicates the cells as represented in (D) for siCCDC66. Skewed cells with asymmetric cleavage furrow are counted, and percentage is calculated according to total telophase/cytokinesis cell number. Data represent the mean ±SEM of three independent experiments. (**p<0.01). (F) *In vitro* microtubule bundling with full length CCDC66. His-MBP-mNG-CCDC66 was purified from insect cells using Ni-NTA agarose beads and microtubule bundling was performed. Alpha-tubulin was visualized by rhodamine labelled tubulin and CCDC66 was visualized with mNG signal. Scale bar: 5 µm. (G) *In vitro* microtubule bundling with CCDC66 (570-948). His-MBG-CCDC66 (570-948) C terminal fragment was purified from bacterial culture using Ni-NTA agarose beads and microtubule bundling was performed. Alpha tubulin was visualized by rhodamine labelled tubulin. Scale bar: 5 µm. (H) Spatial distribution of PRC1 at the spindle midzone upon CCDC66 depletion. U2OS cells were transfected with the indicated siRNAs, fixed with methanol and stained for alpha-tubulin, PRC1 and DAPI. Images represent anaphase cells, which are the colored versions of the inverted images presented in (A). Scale bar: 5 µm. The graph represents the plot profile of PRC1 and alpha tubulin. Using ImageJ, a straight line (thickness 200) was drawn pole-to-pole direction covering the PRC1 area and intensity along the distance was plotted on Graphpad Prism. Model shows representation of how the plot profile was generated.

The width of the midzone remained unaltered upon CCDC66 depletion (Fig. S5A). In addition to the central spindle, CCDC66 loss compromised the geometry of the cleavage furrow and resulted in highly asymmetric cleavage furrow ingression (Fig. 5D). About 63.8% of CCDC66-depleted cells exhibited this defect relative to 23.8% in control cells (Fig. 5E). Taken together, these results demosntrate that CCDC66 is involved in the assembly and organization of central spindle and intercellular bridge MTs.

Defects in central spindle and intercellular bridge/midbody, as well as K-fibers could be attributed to MT bundling activity of CCDC66. To test this, we used *in vitro* experiments to investigate the possible MT crosslinking activity of CCDC66. We expressed MBP-mNG-CCDC66 in insect cells and purified it using nickel beads (Fig. S5C, D). As a control, we purified MBP-mNG in bacteria (Fig. S5F). Using *in vitro* MT sedimentation assay, we showed that MBP-mNG-CCDC66 directly binds to MTs (Fig. S5E). After validation of MT affinity, we performed *in vitro* MT bundling assays. Incubation of MBP-mNG-CCDC66, but not MBP-mNG, with taxol-stabilized rhodamine-labeled MTs resulted in the formation of MT bundles *in vitro* (Fig. 5F). Notably, these bundles co-localized with CCDC66, confirming its direct MT affinity. Given that the C-terminal 570-948 residues of CCDC66 binds to MTs and localizes to spindle MTs, we next asked whether this fragment bundles MTs *in vitro*. We expressed and purified MBP-CCDC66 (570-948) in bacteria (Fig. S5G). Like full length CCDC66, MBP-CCDC66 (570-948) associated with MTs directly and promoted MT bundling *in vitro* (Fig. 5G, S5H). In agreement with their *in vitro* activities, overexpression of mNG fusions of CCDC66 and its C-terminal (570-948) fragment induced formation of MT bundles (Fig. S5I) in cells. Collectively, these results demonstrate that CCDC66 is a cross-linking MAP that is required for assembly and organization of central spindle and midbody MTs during cytokinesis.

MT-binding and bundling activities of PRC1 are required for microtubule organization at the spindle midzone in anaphase, localization of MAPs within this structure and successful completion of cytokinesis (Gruneberg et al., 2006; Jiang et al., 1998; Mollinari et al., 2002; Zhu and Jiang, 2005). Given that CCDC66 interacts and co-localizes with PRC1 (Fig. 2D, 2F), we tested whether the MT disorganization at the spindle midzone and cleavage furrow are due to defective PRC1 targeting to the central spindle and midbody. As revealed by the plot profile analysis of PRC1 intensity, the spatial distribution of PRC1 at the central spindle were disrupted. Specifically, PRC1 signal was spread over a broader region of anaphase B spindle in CCDC66-depleted cells (Fig. 5H). Of note, the fluorescence intensity of PRC1 at the midbody was comparable between control and CCDC66-depleted cells (Fig. S5B). Taken together, these results suggest potential involvement of CCDC66 in regulating recruitment of central spindle components including but not limited to PRC1.

### CCDC66 is required for centrosome maturation and MT nucleation during cell division

One of the most prominent phenotypes associated with CCDC66 depletion is the significant decrease in astral, metaphase spindle and central spindle MT intensities. This finding led us to investigate the functions of CCDC66 during MT nucleation in dividing cells. To this end, we performed MT regrowth experiments in control and CCDC66-depleted cells synchronized using the Eg5-inhibitor S-trityl-L-cysteine (STLC) (Fig. 6A). Following MT depolymerization by nocodazole treatment and its washout, cells were then fixed and stained for MTs and the centriole marker Centrin 3 at the indicated time points (Fig. 6A). The centrosomal and non-centrosomal MT aster size was reduced in CCDC66-depleted cells relative to control cells at 3, 5 and 8 min after washout, which indicates MT nucleation defects (Fig. 6A, 6B). Notably, CCDC66-depleted cells had an increased number of MT nucleating centers than control cells, suggesting possible activation of non-centrosomal MT nucleation pathways (Fig. S6A). CCDC66-depleted cells were also delayed in the formation of the bipolar spindle after nocodazole treatment (Fig. 6C, 6D). By 40 min, 30.9 ± 2.6% control cells and 9.9 ± 0.2% CCDC66-depleted cells formed a bipolar spindle (Fig. 6C, 6D). Supporting its roles in centrosome-mediated and non-centrosomal MT nucleation, gamma-tubulin levels at the spindle poles and microtubules were reduced in CCDC66-depleted cells relative to control cells (Fig. 6E). Notably, ultrastructure expansion microscopy (U-ExM) analysis of STLC-synchronized G2 and mitotic cells confirmed reduced gamma-tubulin recruitment to the PCM and spindle MTs upon CCDC66 depletion (Fig. S6C). We also noted that organization of gamma-tubulin at the PCM was disrupted while its pool at the centriole wall and lumen remained intact(Fig. S6C), which was recently reported to be required for centriole integrity and cilium assembly (Schweizer et al., 2021).

**Figure 6.**
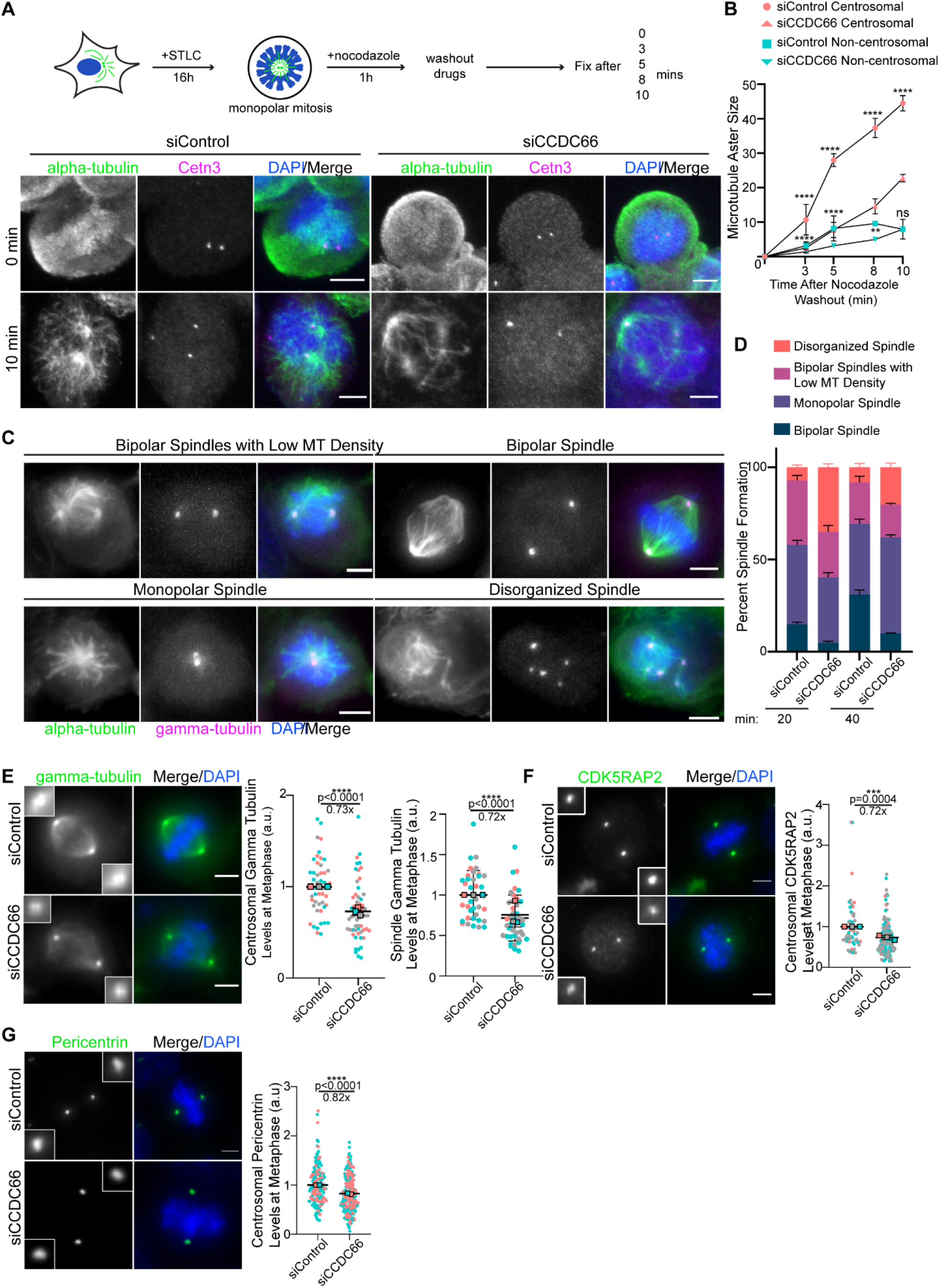
CCDC66 recruits core PCM proteins to the centrosome and is required for mitotic microtubule nucleation. (A) CCDC66 depletion slows down microtubule nucleation in mitotic cells. As illustrated in the experimental plan, U2OS cells were transfected with control and CCDC66 siRNA. 48 h post-transfection, they were synchronized with 5 µM STLC treatment for 16h. After cell synchronization, microtubules were depolymerized by nocodazole treatment for 1h. Following nocodazole wash out, cells were fixed and stained for alpha-tubulin, Centrin 3 and DAPI at the indicated time points. (B) Quantification of A. For quantification of microtubule aster size, centrosomal and acentrosomal microtubule nucleation area is measured on ImageJ using polygon selection tool. Data represent the mean ±SEM of two independent experiments. (****p<0.0001, **p<0.01). Scale bar: 5 µm. (C) Effect of CCDC66 depletion on bipolar spindle assembly. Control and CCDC66 siRNA-transfected cells were stained with alpha-tubulin, gamma-tubulin and DAPI. (D)Quantification of C. Mitotic cells were scored based on their spindle architecture as bipolar spindle, monopolar spindle, bipolar spindles with low microtubule density, and disorganized spindle. Data represent the mean ±SEM of two independent experiments. Scale bar: 5 µm. (E-G) Effects of CCDC66 depletion on abundance of PCM proteins at the spindle poles. U2OS cells were transfected with control and CCDC66 siRNA. After 48 h, cells were fixed with methanol and stained for (C) gamma-tubulin, (D) CDK5RAP2, (E) Pericentrin, and DAPI. Centrosomal abundance of PCM proteins and spindle abundance of gama tubulin was measured on ImageJ by drawing a 3.4 µm^2^ circular area. Data represent mean ±SEM of two (Pericentrin) and three (gamma-tubulin, CDK5RAP2) independent experiments. (****p<0.0001). Images for each panel represent cells captured with the same camera settings from the same coverslip. Scale bar: 5 µm.

During centrosome maturation, centrosomes recruit more PCM proteins to increase their MT-nucleating capacity. CDK5RAP2, CEP192, CEP152 and Pericentrin are required for spindle pole recruitment of gamma-tubulin (Luders, 2012; Palazzo et al., 2000). Given that CCDC66 interacts with these proteins, we tested their centrosomal targeting as a potential mechanism by which CCDC66 regulates centrosome maturation. To test this, we quantified centrosomal levels of these PCM proteins in control and CCDC66 siRNA-transfected cells. The levels of CDK5RAP2 and Pericentrin, but not CEP192 and CEP152, were significantly reduced at the spindle poles in CCDC66-depleted cells as compared to control cells (Fig. 6F, 6G, S6D, S6E). Immunoblot analysis of lysates from control and CCDC66-depleted cells with antibodies against these proteins indicated that CCDC66 loss does not alter their cellular abundance (Fig. S6B). Collectively, these results demonstrate that CCDC66 functions during centrosomal and non-centrosomal MT nucleation via targeting gamma-tubulin to spindle poles and microtubules.

### Expression of CCDC66 and its centrosome and MT-binding fragments restore mitotic and cytokinetic defects in CCDC66-depleted cells to different extents

The functional significance of the dynamic localization of CCDC66 to the spindle poles, and various MT arrays as well as the relative contribution of its MT nucleation and organization activities to its functions are not known known. To distinguish between the function of these different CCDC66 pools and activities during cell division, we performed phenotypic rescue experiments with three different CCDC66 siRNA-resistant constructs: mNG-CCDC66 to validate the specificity of the phenotypes, 2) mNG-CCDC66 (570-948) to assess the functional significance of CCDC66 localization to the spindle poles and MT-binding and bundling activity, 3) C-terminal mNG-CCDC66 fusion with the centrosomal targeting domain PACT to distinguish its centrosome-specific activities from the ones mediated by MTs. Next, we used lentiviral transduction to generate U2OS cells stably expressing these fusion proteins as well as mNG itself as a control and validated their expression at the expected size, siRNA resistance and localization in CCDC66-depleted cells by immunoblotting and immunofluorescence (Fig. 7A, S7A, S7B). In control and CCDC66 siRNA-transfected cells, mitotic localization profiles of mNG-CCDC66 and mNG-CCDC66 (570-948) were similar to the ones we reported in Fig. 1 (Fig. 7A). As for mNG-CCDC66-PACT, its localization was restricted to the centrosome and did not localize to the MTs (Fig. 7A). Of note, its centrosomal to cytoplasmic relative fluorescent intensity was much higher, indicating that the majority of cellular CCDC66 was sequestered at the centrosome (Fig. 7A).

**Figure 7.**
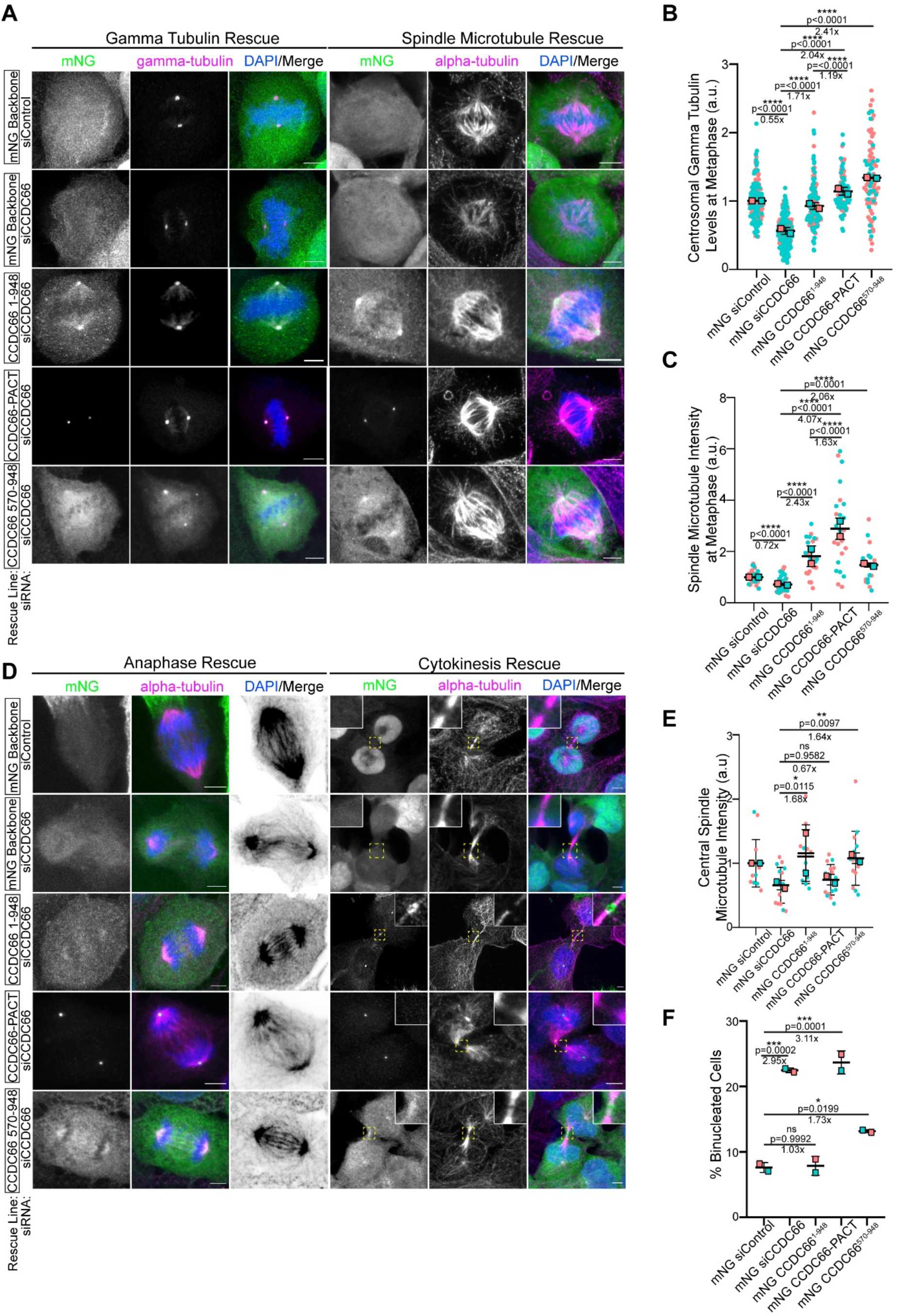
Centrosome and microtubule-affinity of CCDC66 is required for its mitotic and cytokinetic functions to different extents. (A) Representative images for the gamma tubulin and spindle microtubule density rescue experiments performed using U2OS::mNeonGreen, U2OS::mNeonGreen-CCDC66^1-948^ (full-length), U2OS::mNeonGreen-CCDC66-PACT and U2OS::mNeonGreen-CCDC66^570-948^ stable cells. Cells were transfected with control and CCDC66 siRNA. 48 h post-transfection, they were fixed with 4% PFA and stained for either gamma-tubulin or alpha-tubulin and DAPI. Scale bar: 5 µm. (B) Quantification of (A). gamma-tubulin intensity was measured on ImageJ by drawing a 3.4 µm^2^ circular area. Data represent the mean ±SEM of two independent experiments. (****p<0.0001) (C) Quantification of (A). Spindle microtubule intensity was measured on ImageJ by taking several points on the spindle to measure the intensity then taking the average. Data represent the mean ±SEM of two independent experiments. (****p<0.0001). (D) Representative images for the anaphase and cytokinesis rescue experiments performed using U2OS::mNeonGreen, U2OS::mNeonGreen-CCDC66^1-948^, U2OS::mNeonGreen-CCDC66-PACT and U2OS::mNeonGreen-CCDC66^570-948^ stable cells. Cells were transfected with control and CCDC66 siRNA. 48 h post-transfection, they were fixed with 4% PFA and stained for alpha-tubulin and DAPI. Scale bar: 5 µm. (E) Quantification of (D). Graph represents the microtubule density at central spindle. Central spindle microtubule intensity was measured on ImageJ by taking several points on the spindle to measure the intensity and subtracting the background mean intensity. Data represent the mean ±SEM of two independent experiments. (ns: not significant *p<0.05 **p<0.01). (F) Quantification of (D). Graph represents the percentage of binucleated cell number. Data represent the mean ±SEM of two independent experiments. (ns: not significant *p<0.05 ***p<0.005).

Next, we examined whether expression of mNG-fusions of CCDC66, CCDC66 (570-948) and CCDC66-PACT restores defective targeting of gamma-tubulin to the spindle poles, reduced spindle MT intensity in metaphase and anaphase cells, increased cold-sensitivity of K-fibers, spindle mispositioning, disorganized central spindle and increased binucleation in CCDC66-depleted cells. mNG-CCDC66 expression rescued all seven phenotypes to comparable or greater levels to control siRNA-transfected mNG-expressing cells, indicating that these phenotypes are specific to CCDC66 depletion (Fig. 7A-F, S7C-F). Similarly, mNG-CCDC66 (570-948) expression partially or fully rescued all seven phenotypes, suggesting that centrosomal and MT affinity is sufficient for CCDC66 functions in these processes (Fig. 7A-F). As for the spindle MT levels, not only CCDC66 (570-948) but also full length CCDC66 resulted in higher averages relative to control cells, suggesting that their expression might promote these phenotypes via increased MT nucleation. Strikingly, mNG-CCDC66-PACT only restored gamma-tubulin levels at the spindle poles, spindle MT levels and spindle mispositioning defects to comparable or higher levels than that of control cells (Fig. 7A-C). However, it did not rescue defects in K-fiber stability, central spindle MT intensities and organization and cytokinesis, suggesting that the MT-binding and bundling activities of CCDC66 is required for CCDC66 functions at the k-fibers, central spindle and cleavage furrow (Fig. 7D-F, S7D-F). Collectively, these results show that CCDC66 functions during mitosis and cytokinesis via regulating centrosomal and non-centrosomal MT nucleation as well as MT organization. Importantly, the relative contribution of these activities to different CCDC66 functions varies based on the mechanisms by which different MT arrays are assembled and organized.

## Discussion

In this study, we identified the centrosomal and ciliary MAP, CCDC66, as a new player of the machinery governing the assembly and organization of the mitotic and cytokinetic MT arrays and thereby cell cycle progression. As summarized in the model shown in Fig. 8, our findings reveal two important roles of CCDC66 during cell division. First, CCDC66 is required for mitotic progression via regulation of spindle assembly, organization and positioning, levels of spindle MTs, k-fiber integrity and chromosome alignment. Second, CCDC66 functions during cytokinesis in part by regulating assembly and organization of central spindle and midbody MTs. Our work provides new insights into the spatiotemporal regulation of the mitotic and cytokinetic events governed by the dynamic changes of the MT cytoskeleton and centrosomes.

**Figure 8.**
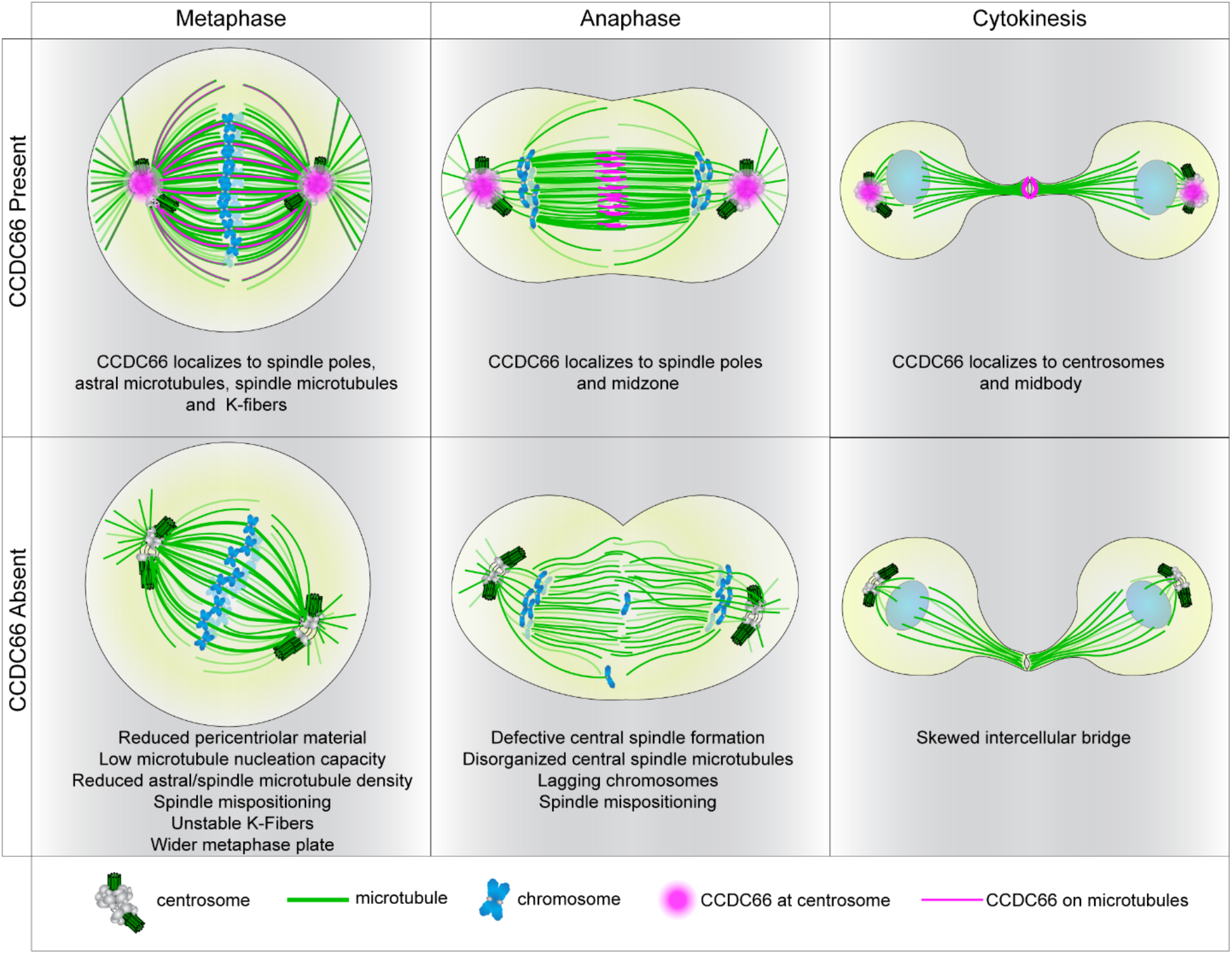
Model CCDC66 localization and functions during mitosis and cytokinesis. The model shows CCDC66 localization during cell division and the phenotypic consequences of CCDC66 depletion on mitotic and cytokinetic MT arrays as well as chromosomes. In metaphase cells, CCDC66 (magenta) localizes to the spindle poles, the astral microtubules, the spindle microtubules and K-fibers. CCDC66 loss results in a reduced pericentriolar material size, microtubule nucleation capacity, astral/spindle microtubule density as well as unstable K-fibers. It also causes a shift in orientation and increases metaphase plate width. During anaphase, CCDC66 localizes to the spindle poles and midzone. CCDC66 depletion causes defective central spindle formation, reduction in central spindle microtubule intensity, lagging chromosomes and orientation shift. During cytokinesis, CCDC66 additionally localizes to the centrosomes and the midbody. CCDC66 depletion causes asymmetric cleavage furrow formation

The results of our study revealed MT nucleation and organization as the two major mechanisms by which CCDC66 functions during cell division. In mitotic cells, CCDC66 regulates centrosome maturation via recruitment of core PCM proteins and is required for MT nucleation from the centrosomes during bipolar spindle assembly and positioning. The inability of centrosome-restricted mNG-CCDC66-PACT to rescue central spindle, cytokinesis and k-fiber defects show that CCDC66 does not regulate these processes via centrosomal MT nucleation. Furthermore, two lines of evidence indicate that CCDC66 is also involved in MT-mediated MT nucleation during spindle assembly. First, depletion of CCDC66 resulted in a decrease in microtubule nucleation from non-centrosomal nucleation centers in mitotic cells as well as in gamma-tubulin recruitment to the spindle. Second, CCDC66 depletion resulted in decreased MT density of central spindle in anaphase, which represents *de novo* non-centrosomal MT generation in inter-chromosomal region and requires MT-associated proteins that participate in nucleation, stabilization and bundling (Courtheoux et al., 2019; Uehara and Goshima, 2010). The role of CCDC66 in MT-mediated MT nucleation is further supported by its proximity interactions with all 8 subunits of the HAUS/Augmin complex, which recruits γ-TuRC to MTs and promotes nucleation from spindle MTs (Gheiratmand et al., 2019). Future studies are required to investigate whether and if so how CCDC66 regulates HAUS complex-dependent MT nucleation. It will be worthwhile to characterize direct contribution of CCDC66 on gamma and alpha/beta-tubulin recruitment and *de novo* MT formation *in vitro* in future studies.

The second mechanism by which CCDC66 operates during mitosis is organization of MTs via its crosslinking activity. Using *in vitro* and cellular experiments, we showed that CCDC66 bundles MTs in part via its C-terminal MT-binding domain. In line with our results, a recent study published during the revision of our manuscript reported that GFP-CCDC66 purified from mammalian cells binds to MTs and bundles them *in vitro* (Jijumon et al., 2022). Bundling and stabilization of MTs is essential for the assembly and maintenance of the MT arrays such as the k-fibers, central spindle and midbody (Glotzer, 2009). Consistent with its *in vitro* MT bundling activity, CCDC66 depletion resulted in disorganized MTs at the k-fibers, central spindle and cleavage furrow and these defects were rescued by expression of mNG-CCDC66 (570-948), but not mNGCCDC66-PACT. While these results identify CCDC66 as a new player of organization of stable MT bundles during cell division, the mechanisms by which CCDC66 bundles MTs and the nature of these bundles are not known. Notably, CCDC66 interacts and co-localizes with PRC1 at the centrals spindle and midbody and its depletion disrupts PRC1 distribution at the central spindle, suggesting that CCDC66 regulates cytokinesis in part via PRC1. Further studies aimed at addressing how CCDC66 works together with PRC1 and other components of the central spindle and midbody, as well as with the other factors contributing to contractile actomyosin ring (AMR) formation, will be critical in providing mechanistic insight into CCDC66 functions during cytokinesis.

Despite its roles in MT nucleation and organization, CCDC66 depletion did not result in shorter spindles, which can be explained by a compensatory mechanism activated in CCDC66-depleted cells. Spatiotemporal regulation of MT polymerization, depolymerization, and sliding is critical to spindle length maintenance, providing remarkable ability of metaphase spindles to correct transient fluctuations in morphology (Dumont and Mitchison, 2009; Goshima et al., 2005). Other MAPs and molecular motors that regulate MT stability, dynamics, sliding, as well as regulators of chromatid cohesion and chromosome MT nucleation, probably compensate and correct MT perturbation in CCDC66 absence to maintain steady-state spindle length (Bird and Hyman, 2008). Further characterization of the functional relationship of CCDC66 with the known mitotic MAPs and *in vitro* MT reconstitution assays will contribute to better understanding of regulation of spindle properties by CCDC66.

The pleiotropic localization and activities of CCDC66 during cell division presents challenges in specifically defining mechanisms that underlie its mitotic and cytokinetic functions. Are they regulated by centrosomal and microtubule-associated pools of CCDC66 or their coordinated activity? While we first aimed to address this via identification of CCDC66 fragments that contribute to a single localization and activity, we could not identify such regions. As an alternative, we performed phenotypic rescue experiments with CCDC66 (570-948), which bundles MTs and localizes to centrosomes and CCDC66-PACT, which exclusively localizes to the centrosome. Of note, the strong centrosome affinity of the PACT domain increased CCDC66 levels at the centrosome and created more binding sites for its interactors such as gamma-tubulin, which might have compensated for lack of MT association for a subset of CCDC66 functions. Defective spindle MT density, recruitment of gamma-tubulin to the spindle poles and spindle positioning were rescued both by CCDC66-PACT and CCDC66 (570-948), albeit to different extents. These results suggest that centrosomal and MT pools cooperate in these processes. Strikingly, k-fiber, central spindle and cytokinesis defects were rescued only by CCDC66 (570-948). Given the essential role of chromosome and MT-dependent MT nucleation and bundled MT arrays of the central spindle and midbody during cytokinesis, these results suggest that CCDC66-mediated MT bundling and non-centrosomal MT nucleation are required for these cellular processes (Uehara and Goshima, 2010; Uehara et al., 2009).

CCDC66 has been implicated in several developmental disorders including retinal degeneration and Joubert syndrome (Dekomien et al., 2010; Gerding et al., 2011; Latour et al., 2020; Murgiano et al., 2020; Schreiber et al., 2018). Consistent with its link to ciliopathies, we and others previously showed that retinal degeneration mutations disrupt its ciliary functions and interactions (Conkar et al., 2017; Murgiano et al., 2020). Importantly, our results suggest that disruption of its nonciliary functions of CCDC66 might also contribute to disease pathogenesis. For example, CCDC66 is required for proper spindle positioning, which is critical for specification of the site of the cleavage furrow and distribution of cell fate determinants to daughter cells during architecture and organization of tissues affected in ciliopathies (Kotak, 2019; Kotak and Gonczy, 2013). We note that CCDC66^-/-^ mice were embryonically viable and did not develop tumors, indicating that defective cell cycle defects linked to CCDC66 loss alone do not disrupt embryogenesis and is not tumorigenic

## Materials and Methods Plasmids

pDEST-GFP-CCDC66 and pDEST-GFP-CCDC66^RR^ plasmids used for transfection and transformation experiments were previously described (Conkar et al., 2017). Full-length CCDC66, CCDC66 (570-948) and CCDC66-PACT was amplified by PCR and cloned into pCDH-EF1-mNeonGreen-T2A-Puro lentiviral expression plasmid and pcDNA5.1-FRT/TO-FLAG-miniTurbo mammalian expression plasmid. CCDC66 (570-948) was cloned into pDEST-His-MBP expression plasmid using Gateway cloning (Thermo Scientific). siRNA resistant mNG-CCDC66 was amplified from siRNA resistant GFP-CCDC66^RR^ plasmid and cloned into pCDH-EF1-mNeonGreen-T2A-Puro plasmid. EB3-mNeonGreen-T2A-gamma-tubulin-tagRFP plasmid was a gift from Andrew Holland (Johns Hopkins University School of Medicine) (Yeow et al., 2020). Myc-BirA* fusions of CDK5RAP2, CEP192 and CEP152 were previously described (Firat-Karalar et al., 2014). PLK1 was amplified by PCR and cloned into pDEST-Myc-BirA* plasmid using Gateway cloning. GFP-CEP55 plasmid was a gift from Kerstin Kutsche (UKE, Hamburg). GFP-PRC1 plasmid was a gift from Xuebiao Yao (Morehouse School of Medicine). (Gerding et al., 2011).This might be due to the compensatory mechanisms activated upon chronic loss of CCDC66, which is supported by its evolutionary conservation profile. CCDC66 is not as highly conserved as the HAUS complex or the core PCM proteins that it interacts with, suggesting that it is not an essential conserved player of cell division, but instead vertebrate-specific regulatory protein required for regulating the fidelity of cell division. Future studies are required to determine whether nonciliary functions of CCDC66 contribute to developmental disorders.

mNeonGreen and mNeonGreen-CCDC66 were amplified by PCR and coned into pDONR221 using Gateway recombination. Subsequent Gateway recombination reactions using pDEST-His-MBP (gift from David Waugh - Addgene plasmid # 11085) and pFastBac-DEST (gift from Tim Stearns; Gateway R1R2 destination cassette was cloned into pFastBac-HT-MBP-D using KpnI and XhoI) were performed to generate His-MBP-mNeonGreen and His-MBP-mNeonGreen-CCDC66 for expression for expression in bacterial cells and insect cells, respectively.

### Cell culture and transfection

Human telomerase immortalized retinal pigment epithelium cells (hTERT-RPE1, ATCC, CRL-4000) were cultured with Dulbecco’s modified Eagle’s Medium DMEM/F12 50/50 medium (Pan Biotech, Cat. # P04-41250) supplemented with 10% Fetal Bovine Serum (FBS, Life Technologies, Ref. # 10270-106, Lot # 42Q5283K) and1% penicillin-streptomycin (GIBCO, Cat. # 1540-122). Human embryonic kidney (HEK293T, ATCC, CRL-3216), and osteosarcoma epithelial (U2OS, ATCC, HTB-96) cells were cultured with DMEM medium (Pan Biotech, Cat. # P04-03590) supplemented with10% FBS and 1% penicillin-streptomycin. All cell lines were authenticated by Multiplex Cell Line Authentication (MCA) and were tested for mycoplasma by MycoAlert Mycoplasma Detection Kit (Lonza). U2OS cells were transfected with the plasmids using Lipofectamine 2000 and according to the manufacturer’s instructions (Thermo Scientific Scientific). HEK293T cells were transfected with the plasmids using 1mg/ml polyethylenimine, MW 25 kDa (PEI). For microtubule depolymerization experiments, cells were treated with 5 µg/ml nocodazole (Sigma-Aldrich, Cat. #M1404) or vehicle (dimethyl sulfoxide) for one hour at 37 C. For cell synchronization experiments, 5 µM (+)-S-trityl-L-cysteine (STLC) (Alfa-Aesar, Cat. #2799-07-7) was used for 16 h at 37 C.

### Lentivirus production and cell transduction

Lentivirus were generated using pcDH-mNG-CCDC66, pcDH-mNG-CCDC66 (570-948), pcDH-mNG-CCDC66-PACT and pCDH-EF1-mNeonGreen-T2A-Puro, and pLVPT2-mCherry-H2B plasmids as transfer vectors. For infection, 1 × 10^5^ U2OS cells were seeded on 6-well tissue culture plates the day before infection, which were infected with 1 mL of viral supernatant the following day. 48-hour post-infection, cells were split and selected in the presence of 6mg/ml puromycin for RPE1s and 4mg/ml puromycin for U2OS cells for four to six days until all the control cells died. U2OS and RPE1 cells stably expressing mCherry-H2B and mNeonGreen fusions of full-length CCDC66 and its truncations were generated by infection of cells with lentivirus expressing the fusions.

### siRNA and rescue experiments

CCDC66 was depleted using an siRNA with the sequence 5′-CAGTGTAATCAGTTCACAAtt-3′. Silencer Select Negative Control No. 1 (Thermo Scientific) was used as a control (Conkar et al., 2017). siRNAs were transfected into U2OS cells with Lipofectamine RNAiMax according to the manufacturer’s instructions (Thermo Scientific Scientific). For rescue experiments, U2OS cells stably expressing mNG or mNG–CCDC66, mNG-CCDC66 PACT and mNG-CCDC66 570-948 were transfected with control and CCDC66 siRNAs by using Lipofectamine RNAiMax (Thermo Scientific Scientific). 48 h post transfection with the siRNAs, cells were fixed and stained. Due to the heterogeneity of the expression of fusion proteins, stable cells in which fusion proteins are not overexpressed and localize properly were accounted for quantification of phenotypic defects.

### Protein expression and purification

Protein expression in Rosetta (BL21(DE3)) cells transformed with respective constructs was induced by addition of 0.5 mM IPTG at OD600 of 0.5–0.6 for 16 hr at 18°C. BEVS baculovirus expression system and protocol (Fitzgerald et al., 2006) was used for expression of tagged full length CCDC66 protein. Briefly, 100 mL of Hi5 cells (1 × 106 cell/mL) were infected with P1 baculovirus produced in Sf9 cells, carrying His-MBP-mNeonGreen-CCDC66, at MOI of 1. Cells were collected 48 hr after infection.

For protein purification, cells were lysed by sonication in lysis buffer (20 mM Hepes, pH 7.0 (or pH 7.5 for full length protein), 250 mM NaCl, 0.1% Tween20, 2 mg/ml lysozyme, 1 mM PMSF, 1mM protease inhibitor cocktail, 5mM BME and 10 mM Imidazole) and clarified at 19,000 rpm for 1 hr at 4°C. His-tagged proteins were subsequently purified using Ni-NTA–agarose beads (Thermo Scientific). Proteins were eluted with elution buffer (20 mM Hepes, pH 7.0 or pH7.5, 250 mM NaCl, 5mM BME and 250 mM imidazole). For subsequent microtubule assays, proteins were dialyzed against BRB80 buffer (80 mM PIPES pH 6.8, 1 mM EGTA, 1m M MgCl_2_).

### *In Vitro* MT bundling assay

Fluorescent MTs were polymerized at 2 mg/ml by incubating tubulin and rhodamine labeled tubulin (Cytoskeleton, Inc.) at 10:1 ratio in BRB80 with 1 mM DTT and 1mM GTP for 5 min on ice, then preclearing by centrifuging for 10 min at 90000 rpm at 2°C in TLA100 rotor. Cleared tubulin mixture was polymerized at 37°C by adding taxol and increasing concentration stepwise, with final concentration of 20 µM. MTs were pelleted over warm 40% glycerol BRB80 cushion at 70000 rpm for 20 min. After washes with 0.5% Triton-X100, pellet was resuspended in 80% of the starting volume of warm BRB80 buffer with 1mM DTT and 20 µM Taxol.

Bundling assays were performed as previously described (Tao et al., 2016). Briefly, 100 nM of protein His-MBP-CCDC66570-948, His-MBP-mNeonGreen, His-MBP-mNeonGreen-CCDC66 or MBP was mixed with 2 µM MTs and 20 µM taxol in buffer T (20mM Tris, pH 8.0, 150mM KCl, 2mM MgCl2, 1mM DTT, protease inhibitors) for 30 min rocking at room temperature. The reaction mixtures were transferred into a flow chamber under a HCl-treated coverslip, and unstuck proteins were washed out with the excess of buffer T. Bundling of fluorescent MTs was observed with a Leica SP8 confocal microscope and 63x 1.4 NA oil objective (Leica Mycrosystems). Experiment was repeated 4 times.

### *In Vitro* MT sedimentation assay

Purified bovine brain tubulin (Cytoskeleton Inc.) was precleared at 90000 rpm for 5 minutes at 2°C with TLA100 rotor. Cleared tubulin was polymerized at 1 mg/ml in the presence of 1 mM GTP and increasing concentrations of taxol to a final concentration of 20 µM. After incubation of microtubules at room temperature overnight, 500 nM of purified protein was mixed with 3 µM taxol-stabilized microtubules or BRB80 buffer with taxol and incubated for 30 minutes at room temperature. Samples were loaded onto 60% glycerol BRB80 cushions and centrifuged at 50000 rpm for 30 min with TLA100 rotor at room temperature. Supernatants were collected, glycerol cushions removed by aspiration, and pellets solubilized in SDS sample buffer. Equivalent volumes of supernatant and pellet fractions were resolved by SDS-PAGE.

### Ultra-Structure Expansion Microscopy (U-ExM)

U-ExM was performed as previously described (Gambarotto et al., 2019). Briefly, U2OS cells were transfected with siControl or siCCDC66. 48 hours after transfection, cells were treated with 5 µM STLC (Alfa-Aesar, Cat. #2799-07-7) for 16 h. Coverslips were incubated in 1.4% formaldehyde / 2% acrylamide (2X FA / AA) solution in 1X PBS for 5 h at 37 C prior to gelation in Monomer Solution supplemented with TEMED and APS (final concentration of 0.5%) for 1 hr at 37°C. Denaturation was performed at 95 C for 1h30 and gels were stained with primary antibodies for 3 h at 37 C. Gels were washed 3 × 10 min at RT with 1 X PBS with 0.1% Triton-X (PBST) prior to secondary antibody incubation for 2h30 at 37 C followed by 3 × 10 min washes in PBST at RT. Gels were expanded in 3 × 150 ml dH2O before imaging. The following reagents were used in U-ExM experiment: formaldehyde (FA, 36.5– 38%, F8775, Sigma-Aldrich), acrylamide (AA, 40%, A4058, Sigma-Aldrich), N,N’-methylenbisacrylamide (BIS, 2%, M1533, SIGMA), sodium acrylate (SA, 97–99%, 408220, Sigma-Aldrich), ammonium persulfate (APS, 17874, Thermo Scientific), tetramethylethylendiamine (TEMED, 17919, Thermo Scientific), and poly-D-Lysine (A3890401, Gibco).

### Immunofluorescence, antibodies, and microscopy

Cells were grown on coverslips, washed twice with PBS, and fixed in either ice cold methanol at - 20 °C for 10 minutes or 4% PFA in Cytoskeletal Buffer ((100 mM NaCl (Sigma-Aldrich, S9888), 300 mM sucrose (Sigma-Aldrich, S0389), 3 mM MgCl2 (Sigma-Aldrich, M2670), and 10 mM PIPES (Sigma-Aldrich, P6757). For CCDC66 endogenous staining with the rabbit polyclonal antibody, cells were first fixed with methanol at - 20 °C, then with 100% acetone for 1 min at room temperature. After rehydration in PBS, cells were blocked with 3% BSA (Capricorn Scientific, Cat. # BSA-1T) in PBS followed by incubation with primary antibodies in blocking solution for 1 hour at room temperature. Cells were washed three times with PBS and incubated with secondary antibodies and DAPI (Thermo Scientific, cat#D1306) at1:2000 for 45 minutes at room temperature. Following three washes with PBS, cells were mounted using Mowiol mounting medium containing N-propyl gallate (Sigma-Aldrich). Primary antibodies used for immunofluorescence were rabbit anti-CCDC66 (Bethyl, A303-339A),, mouse anti gamma-tubulin (Sigma, clone GTU-88, T5326) at 1:1000, rabbit anti GFP at 1:2000 (custom made) (Firat-Karalar et al., 2014), mouse anti alpha-tubulin (Sigma, DM1A) at 1:1000, rabbit anti CEP152 (Bethyl, A302-480A) at 1:500, rabbit anti-CEP192 (Proteintech, 18832 1 AP) at 1:1000, rabbit anti-phospho-Histone H3 at 1:1000, rabbit anti-CDK5RAP2 (Proteintech, 20061 1 AP) at 1:1000, rabbit anti-Pericentrin (Abcam, ab4448) at 1:2000, rabbit anti-CSPP1 (Proteintech, 11931 1 AP) at 1:1000, rabbit anti-PRC1 (Proteintech, 15617 1 AP) at 1:1000, rabbit anti-Cep55 (Proteintech, 23891 1 AP) at 1:1000, rabbit anti-Kif23 (Proteintech, 28587 1 AP), rabbit anti-pAurora A/B/C (Cell Signaling Technology, CST #2914) at 1:1000, and mouse anti-mNeonGreen (Chromotek, 32F6) at 1:500, rabbit anti-GFP was generated and used for immunofluorescence as previously described (Firat-Karalar et al., 2014; Hatch et al., 2010). Secondary antibodies used for immunofluorescence experiments were AlexaFluor 488-, 568-or 633-coupled (Life Technologies) and they were used at 1:2000.

Time lapse live imaging was performed with Leica SP8 confocal microscope equipped with an incubation chamber. For cell cycle experiments, asynchronous cells were imaged at 37C with 5% CO2 with a frequency of 6 minutes per frame with 1.5mm step size and 12 mm stack size in 512×512 pixel format at a specific position using HC PL FLUOTAR 20x/0.50 DRY objective. For centrosomal protein level quantifications, images were acquired with Leica DMi8 inverted fluorescent microscope with a stack size of 8mm and step size of 0.3 mm in 1024×1024 format using HC PL APO CS2 63x 1.4 NA oil objective. Higher resolution images were taken by using HC PL APO CS2 63x 1.4 NA oil objective with Leica SP8 confocal microscope.

Quantitative immunofluorescence for CEP192, CEP152, CDK5RAP2, Pericentrin, gamma-Tubulin, PRC1 and Alpha-Tubulin was performed by acquiring a z stack of control and depleted cells using identical gain and exposure settings. The centrosome region for each cell were defined by staining for a centrosomal marker including gamma-tubulin. The region of interest that encompassed the centrosome was defined as a circle 3.4-mm^2^ area centered on the centrosome in each cell. Total pixel intensity of fluorescence within the region of interest was measured using ImageJ (National Institutes of Health, Bethesda, MD). Background subtraction was performed by quantifying fluorescence intensity of a region of equal dimensions in the area proximate to the centrosome. Statistical analysis was done by normalizing these values to their mean.

### Spindle angle, length, and pole width measurements

U2OS cells were grown on coverslips and transfected with control and CCDC66 siRNA, fixed with methanol and stained for gamma-tubulin. Images are acquired with Leica DMi8 inverted fluorescence microscope at 1024×1024 format with 0.3 mm z step size. Spindle angle is calculated by the formula α=180*tan-1(h/L)/π where h represents the z stack difference between two spindle poles, L represents the distance between spindle poles. Spindle length is calculated by the formula SL= √(h2+L2) where h represents the stack difference between two spindle poles, L represents the distance between spindle poles. Spindle pole width is calculated by measuring the length of the pericentriolar material of the spindle pole as described in the illustrations in Fig. 4.

### Quantitative analysis of the spindle and astral microtubule intensity

U2OS cells were grown on coverslips and transfected with control and CCDC66 siRNA, fixed with methanol and stained for alpha-tubulin. Images were acquired with Leica SP8 Confocal microscopy at 1024×1024 format with 2x zoom factor. For quantification of the astral microtubule intensity, 5 ROIs having XY size 2 microns were positioned manually on the astral microtubules and intensity was recorded. Background of the same ROI was measured in cytoplasm and subtracted from the average signal intensity. For astral microtubule length, the longest microtubule was measured using the length tool on ImageJ. For quantification of spindle microtubule intensity, 10 ROIs having XY size 2 microns were positioned manually on the spindle microtubules and intensity was recorded. Background of the same ROI was measured in cytoplasm and subtracted from the average signal intensity. The values of control and CCDC66 siRNA-treated metaphase cells were plotted relative to mean intensity of control siRNA.

### Quantitative analysis of the central spindle microtubule intensity and morphology

U2OS cells were grown on coverslips and transfected with control and CCDC66 siRNA, fixed with methanol and stained for alpha-tubulin. Images were acquired with Leica SP8 Confocal microscopy at 1024×1024 format with 2x zoom factor. For quantification of central spindle microtubule intensity, 10 ROIs having XY size 2 microns were positioned manually on the central spindle microtubules and intensity was recorded. Background of the same ROI was measured in cytoplasm and subtracted from the average signal intensity. Aberrant central spindles are judged from the disorganization of the microtubules.

### Microtubule regrowth assay

U2OS cells were grown on poly-L-lysine coated coverslips and treated with control or CCDC66 siRNA. After 48 h of transfection, cells were treated with 5 µM/ml STLC for 16h. Next day, cells were treated with 5 µg/ml nocodazole for 1 at 37 °C. Cells were washed extensively with cold PBS to prevent MT polymerization then incubated with warm media and fixed at indicated time points and stained for alpha tubulin and centrin. Images were acquired with Leica SP8 Confocal microscopy at 1024×1024 format with 2x zoom factor. To quantify MT nucleation area (aster size), polygon selection tool on ImageJ is used. Centrosomal and non-centrosomal MT nucleation points were defined based on centrin staining, which marks the centrioles.

### Immunoprecipitation

HEK293T cells were co-transfected with indicated plasmids. 48 post-transfection, cells were washed and lysed with lysis buffer (50 mM HEPES pH 8, 100 mM KCl, 2 mM EDTA, 10% glycerol, 0.1 % NP-40, Protease inhibitors 1:100 pro-block PIC + 1:100 PMSF) for 45 min. Lysates were centrifuged at 13000 rpm for 10 min at 4°C, and supernatants were transferred to a tube. 100 µl from each sample was saved as input. The rest of the supernatant was immunoprecipitated with anti-FLAG M2 agarose beads (Sigma-Aldrich) over-night at 4°C. After washing 3X with lysis buffer, samples were resuspended in SDS containing sample buffer and analyzed by immunoblotting.

### Cell lysis and immunoblotting

Cells were lysed in 50 mM Tris (pH 7.6), 150 mM NaCI, 1% Triton X-100 and protease inhibitors for 30 min at 4C followed by centrifugation at 15.000 g for 15 min. The protein concentration of the resulting supernatants was determined with the Bradford solution (Bio-Rad Laboratories, CA, USA). For immunoblotting, equal quantities of cell extracts were resolved on SDS-PAGE gels, transferred onto nitrocellulose membranes, blocked with TBST in 5% milk for 1 hour at room temperature. Blots were incubated with primary antibodies diluted in 5% BSA in TBST overnight at 4C, washed with TBST three times for 5 minutes and blotted with secondary antibodies for 1 hour at room temperature. After washing blots with TBST three times for 5 minutes, they were visualized with the LI-COR Odyssey® Infrared Imaging System and software at 169 mm (LI-COR Biosciences). Primary antibodies used for immunoblotting were mouse anti gamma-tubulin (Sigma, clone GTU-88, T5326) at 1:5000, rabbit anti GFP at 1:10000 (homemade), mouse anti alpha-tubulin (Sigma, DM1A) at 1:5000, rabbit anti CEP152 (Bethyl, A302-480A) at 1:1000, rabbit anti-CEP192 (Proteintech, 18832 1 AP) at 1:1000, rabbit anti-Cdk5Rap2 (Proteintech, 20061 1 AP) at 1:1000, rabbit anti-Pericentrin (Abcam, ab4448) at 1:2000, mouse anti-mNG (Chromotek, 32F6) and mouse anti-CCDC66 (sigma SAB1408484) at 1:500. Secondary antibodies used for western blotting experiments were IRDye680- and IRDye800-coupled and were used at 1:15000 (LI-COR Biosciences). Secondary antibodies used for western blotting experiments were IRDye680- and IRDye800-coupled and were used at 1:15000 (LI-COR Biosciences)

### Quantification and statistical analysis

Data were analyzed and plotted using GrapPad Prism 7 (GraphPad, La Jolla, CA). Results are shown as mean ± standard errors (SEM) of the mean. Number of biological replicates are indicated in the figure legends. Two-tailed unpaired t tests and one-way analysis of variance (ANOVA) were applied to compare the statistical significance of the measurements. For data that does not follow normal distribution, we applied non-parametric Mann Whitney test. Error bars reflect SD. Following key is followed for asterisk place holders for p-values in the figures: *p < 0.05, **p < 0.01, ***p < 0.001, **** p < 0.0001 ns. not significant

## Acknowledgements

We acknowledge the Firat-Karalar lab members for insightful discussions regarding this work and Dila Gulensoy and Ezgi Odabasi for cloning the mNG-CCDC66-PACT construct and Efe Begar for purifying mNG protein. This work was supported by ERC Grant 679140 to ENF, EMBO Installation Grant and Young Investigator Award to ENF, TUBITAK BIDEB 120C148 grant to ENF and Marie Sklodowska-Curie Fellowship to JD.

## Author Contributions

Conceptualization, ENF, UB, JD.; Methodology, ENF, UB, JD.; Investigation UB, JD.; Resources, ENF, UB, JD.; Writing—Original Draft, ENF, UB, JD.; Writing—Review & Editing, ENF, UB, JD.; Visualization, UB; Supervision, ENF, JD.; Project Administration, ENF.; Funding Acquisition, ENF, JD.

## Supplemental Information

**Figure S1.**
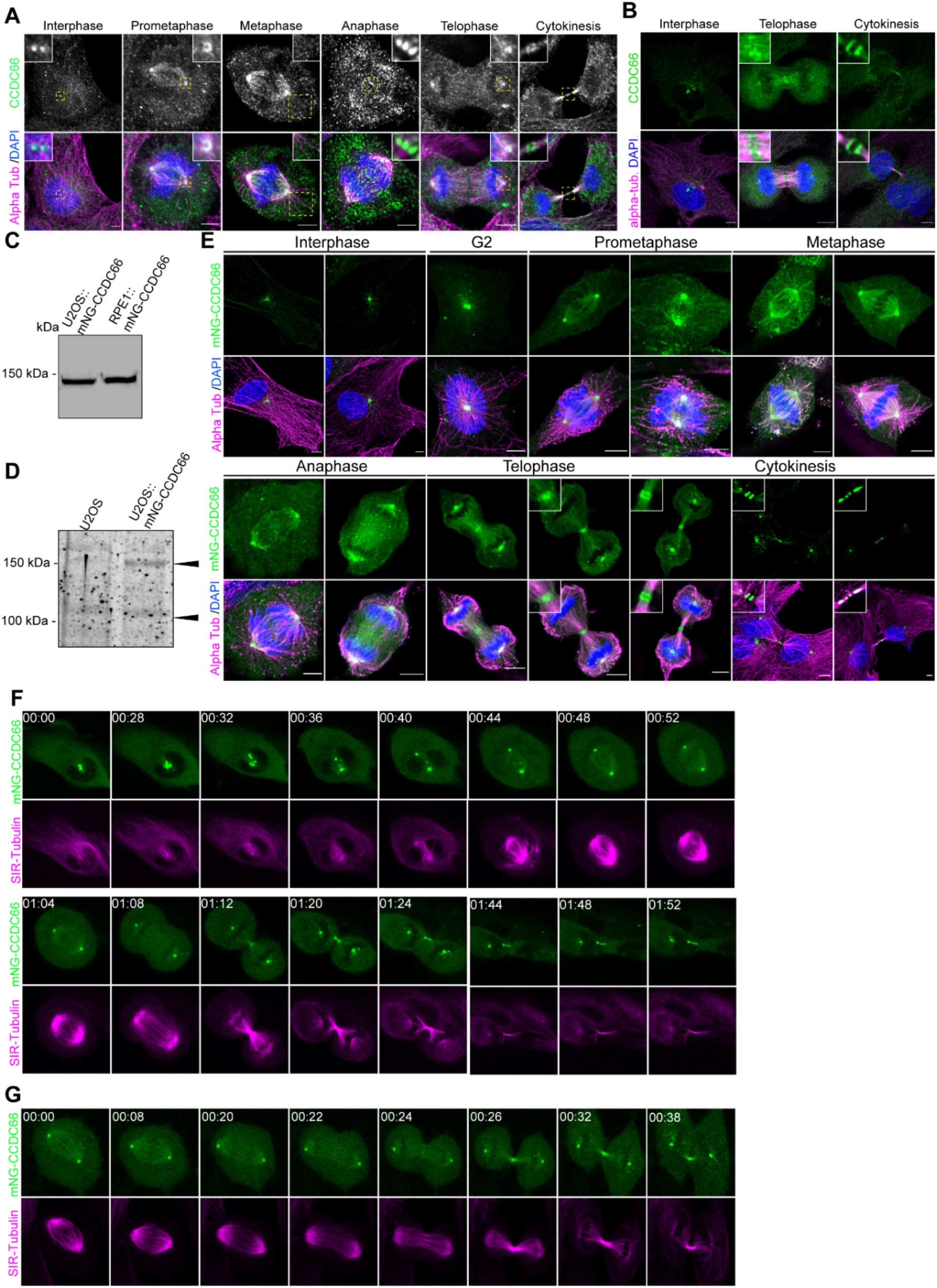
Dynamic CCDC66 localization during cell cycle and validation of mNG-CCDC66-expressing stable cell lines. (A) Localization of CCDC66 at different stages of the cell cycle. U2OS were fixed with methanol followed by acetone and stained for CCDC66, alpha-tubulin and DAPI. Scale bar: 5 µm, insets show 4X magnifications of the boxed regions. (B) Validation of RPE1::mNG-CCDC66 expression with antibody. RPE1::mNG-CCDC66 stable cell line was fixed with MeOH and stained with CCDC66 antibody and alpha-tubulin. Insets show 4X zoom. Scale bar: 5 µm. (C) Validation of mNG-CCDC66 expression in U2OS::mNG-CCDC66 and RPE1::mNG-CCDC66 stable lines by immunoblotting. Extracts from cells were prepared, resolved by SDS-PAGE and blotted with mNG antibody. (D) Relative expression level of mNG-CCDC66 compared to endogenous protein in U2OS cells. Extracts from cells were prepared, resolved by SDS-PAGE and blotted with CCDC66 antibody. (E) Localization of mNeonGreen-CCDC66 at different stages of cell cycle. RPE1 cells stably expressing mNG-CCDC66 fusion (RPE1::mNG-CCDC66) were fixed with 4% PFA and stained for alpha-tubulin and DAPI. Scale bar: 5 µm. (F) Dynamic localization of mNeonGreen-CCDC66 throughout the cell cycle. U2OS cells stably expressing mNeonGreen-CCDC66 fusion (U2OS::mNG-CCDC66) were incubated with 100 nM SiR-Tubulin overnight. Images are acquired every 4 minutes using confocal microscopy. Shown are sixteen time-lapse images from Movie 1 at indicated time points to show dynamic localization of mNG-CCDC66 to spindle poles and microtubules during cell division. (G) Dynamic localization of mNG-CCDC66 throughout the cell cycle. RPE1::mNG-CCDC66 were incubated with 100 nM SiR-Tubulin overnight. Images were acquired every 2 minutes using confocal microscopy. Shown are fourteen time-lapse images from Movie S1 at the indicated time points.

**Figure S2.**
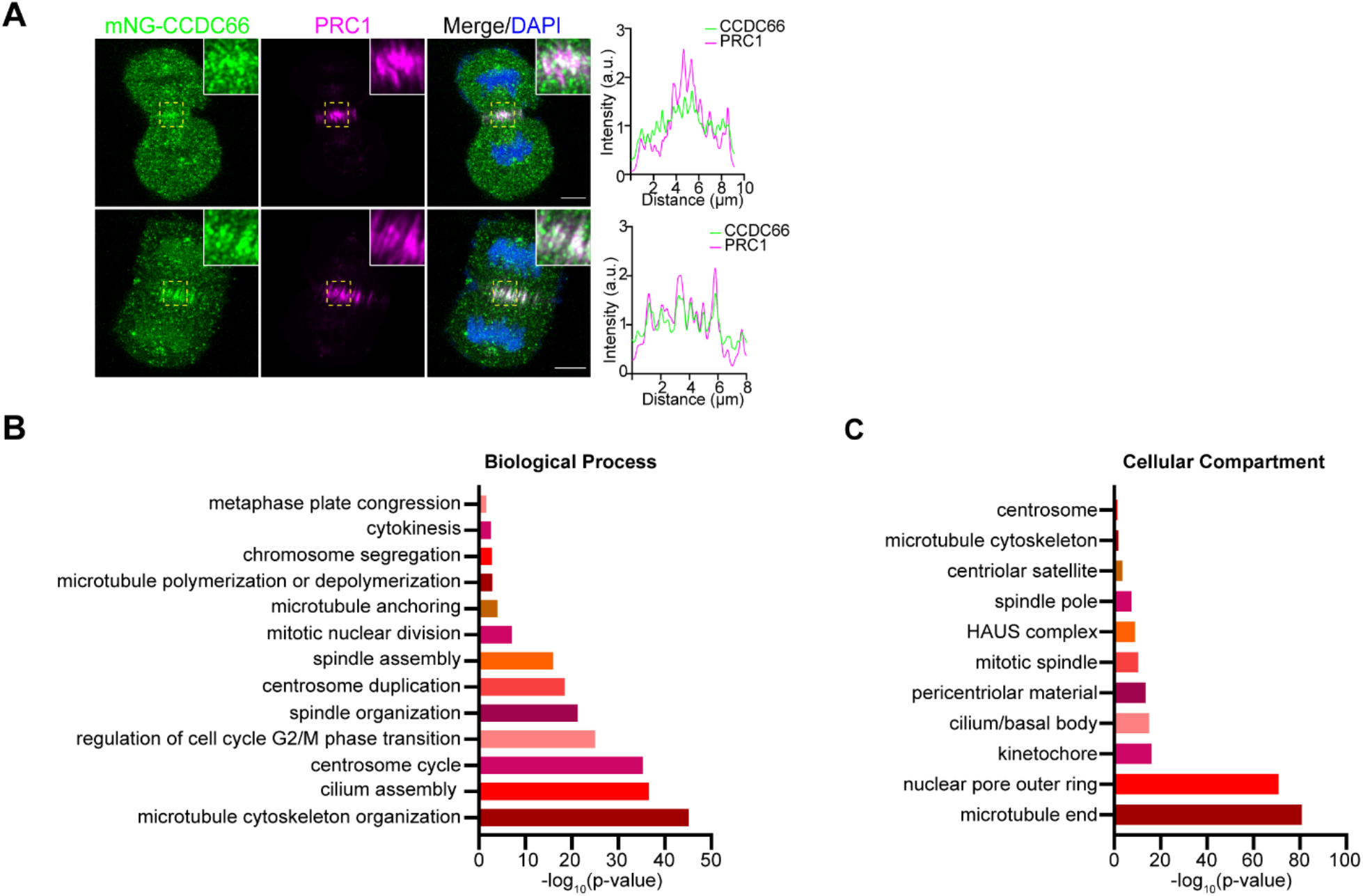
CCDC66 localizes to the central spindle and has extensive proximity interactions with regulators of cells division. (A) Localization of mNG-CCDC66 in RPE1 cells relative to PRC1 during anaphase. RPE1::mNG-CCDC66 cells were fixed with %4 PFA and stained for PRC1 and DAPI. Graphs show the plot profiles to assess co-localization with the indicated marker. Using ImageJ, a straight line was drawn on the midbody and intensity along the distance was plotted on Graphpad Prism. (B,C) GO-enrichment analysis of the CCDC66 proximity interactors based on their (B) biological process and (C) cellular compartment. The x-axis represents the log-transformed p-value (Fisher’s Exact Test) of GO terms.

**Figure S3.**
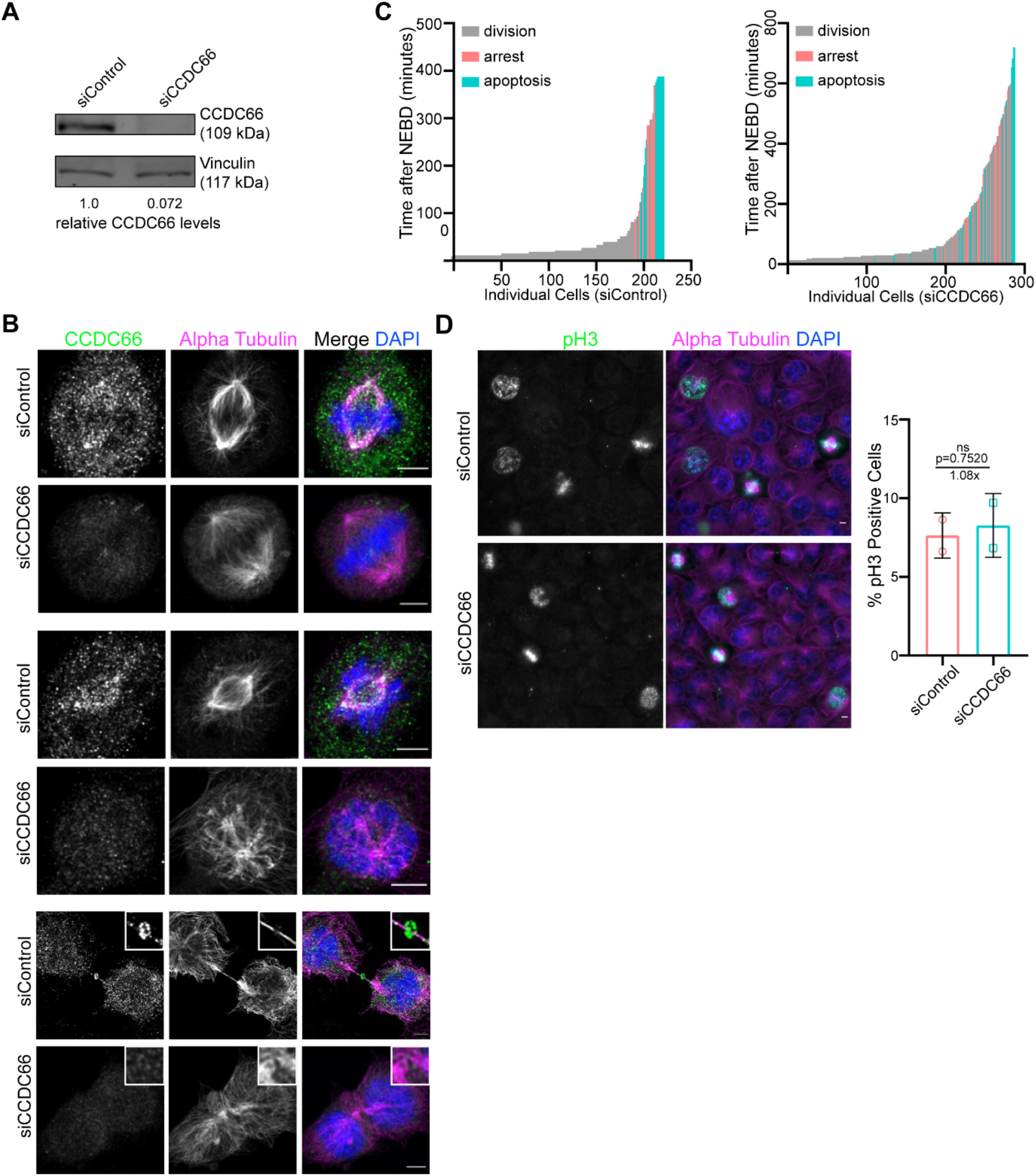
Validation of efficient CCDC66 depletion and its phenotypic consequences on mitotic fate and index. (A) Validation of the efficiency of RNAi-mediated CCDC66 depletion by western blotting and immunofluorescence. U2OS cells were transfected with siControl or siCCDC66. 48 h after post-transfection, cells were fixed and stained with the indicated antibodies. In parallel, cell extracts immunoblotted for CCDC66 and vinculin (loading control). Band intensities were measured on ImageJ and normalized against background and vinculin intensities. Arbitrary value is determined based on siControl. (B) Validation of the CCDC66 antibody by immunofluorescence. U2OS cells were transfected with control or CCDC66 siRNA, fixed with methanol followed by acetone 48 h post-transfection and stained for CCDC66 and alpha tubulin. Representative images are shown at different stages of the cell cycle to indicate the decrease in the signal of CCDC66 upon siCCDC66 transfection. Insets show 4X zoom of boxed areas. Scale bar: 5 µm. (C) Quantification of Figure 3A. The fate of individual cells were plotted as vertical bars, where the height of the bar represents the mitotic time and the color of the bars represent the different fates including successful division (gray), mitotic arrest (pink) and apoptosis (cyan). n>200 cells from each condition was quantified per condition. (D) Effect of CCDC66 depletion on mitotic index. U2OS cells were transfected with control or CCDC66 siRNA, fixed with methanol 48 h post-transfection and stained for the mitotic marker phospho-Histone3 (pH3), alpha-tubulin and DAPI. Mitotic cells are counted based on DNA staining. Data represent the mean ±SEM of two independent experiments. n>1000 for all experiments. Mean mitotic cell number for siControl is 10.31 and mean mitotic cell number for siCCDC66 is 12.49. Representative images are shown. Scale bar: 5 µm.

**Figure S4.**
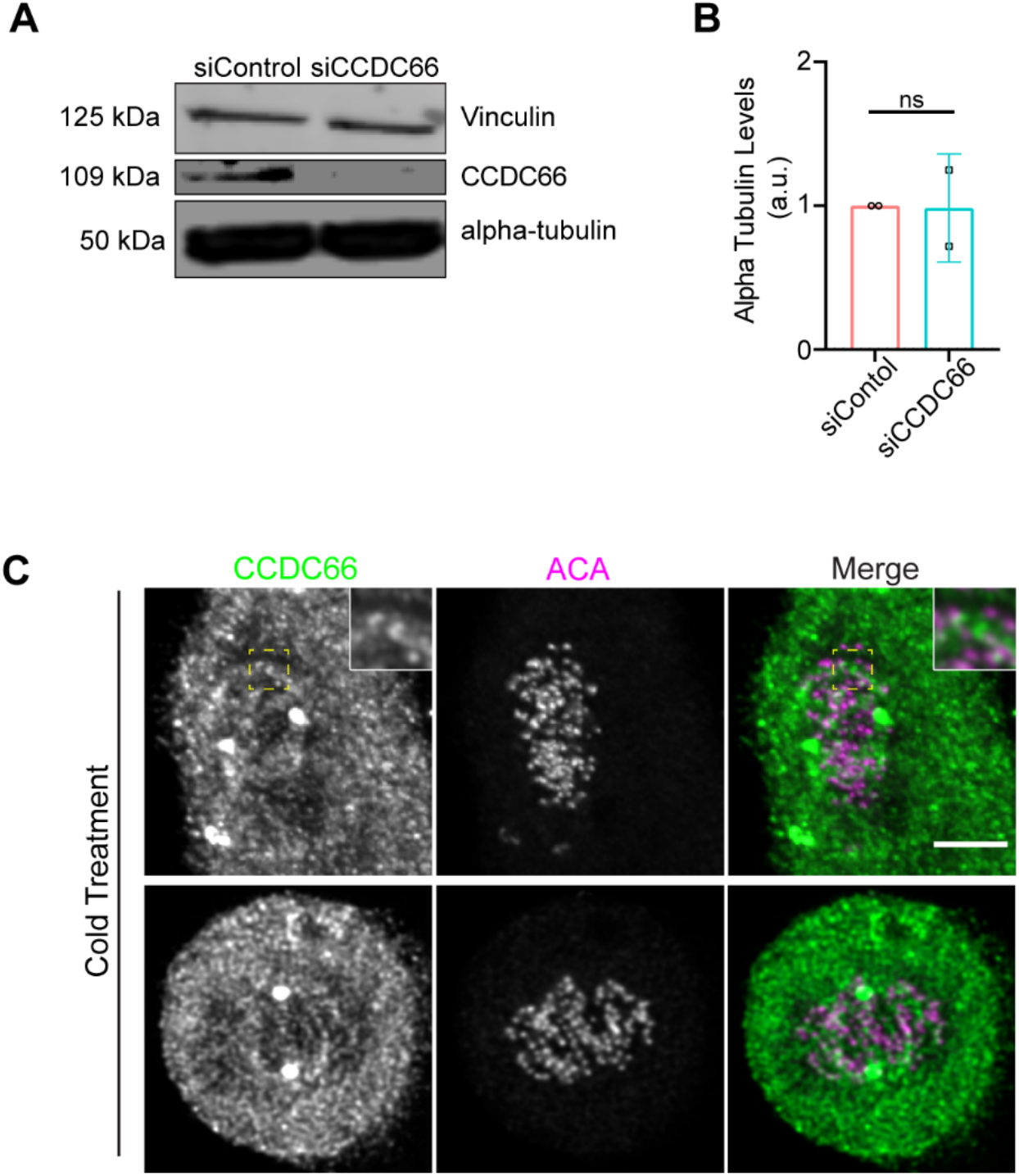
Effects of CCDC66 depletion on cellular abundance of tubulin and endogenous localization of CCDC66 to K-fibers. (A) Cellular abundance of tubulin in control and CCDC66-depleted cells. Cells are transfected with either control or CCDC66 siRNA. After 48 h the lysates are collected and immunoblotted for alpha-tubulin, CCDC66 and vinculin (loading control). (B) Quantification of A. Data represents mean ±SEM of two independent experiments. (ns: not significant). (C) CCDC66 localizes on K-fibers. U2OS cells were grown on coverslips and incubated in ice for 10 minutes before fixation with methanol followed by acetone. Cells were stained for CCDC66, and ACA. Inset shows 4x zoom of boxed area. Scale bar: 5 µm.

**Figure S5.**
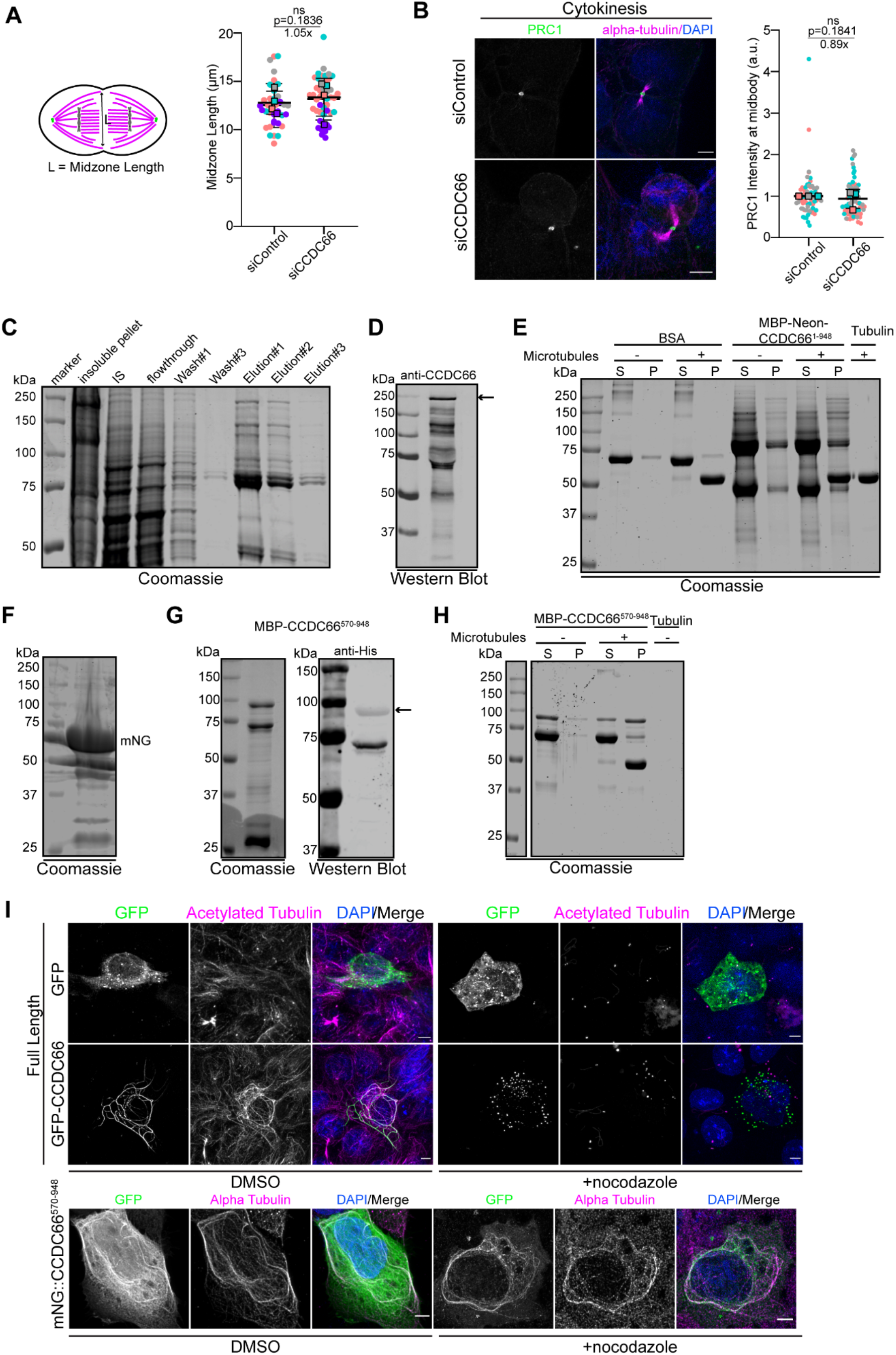
Validation of CCDC66 purification, microtubule association and bundling. (A) Spindle midzone length is not altered by CCDC66 depletion. U2OS cells were transfected with control or CCDC66 siRNA, fixed with methanol followed 48 h post-transfection and stained for alpha tubulin and DAPI. As shown in the representation, midzone length was measured as the distance between the most distant microtubule ends in the midzone. Data represent the mean ±SEM of four independent experiments. (ns: not significant) (B) CCDC66 depletion does not alter PRC1 midbody levels. U2OS cells were transfected with siRNA then fixed with methanol after 48 h and stained for PRC1, alpha-tubulin and DAPI. Images represent cells from Anaphase and cytokinesis. Scale bar: 5 µm. For quantification, PRC1 intensity was measured on ImageJ, the background signal was subtracted, and normalized value was multiplied with area. Arbitrary value was determined by normalizing against siControl. (ns: not significant). (C) His-MBP-mNG-CCDC66 purification. His-MBP-mNG-CCDC66 was purified from insect cells using Ni-NTA agarose beads. Coomassie staining shows the proteins in pellet, nitial sample, flowthrough, wash and elutions. (D) Validation of His-MBP-mNG-CCDC66 purification. Purified His-MBP-mNG-CCDC66 purification was run on SDS-PAGE and blotted with CCDC66 antibody. Arrow corresponds to the full length His-MBP-mNG-CCDC66. (E) His-MBP-mNG-CCDC66 directly interacts with microtubules. His-MBP-mNG-CCDC66 was purified from insect cells and *in vitro* microtubule pelleting was performed and visualized by Coomassie staining. BSA was used as negative control. S stands for supernatant, P stands for pellet. (F) Validation of MBP-mNG purification with Coomassie. (G) Validation of MBP-CCDC66 (570-948) purification with Coomassie. MBP-His-CCDC66 (570-948) was purified from bacterial culture using Ni-NTA agarose beads. Purified protein was run on SDS-PAGE. Coomassie staining and western blotting with anti-His antibody shows the purified protein. (H) MBP-CCDC66 (570-948) directly interacts with microtubules. MBP-His-CCDC66 (570-948) was purified from bacterial culture using Ni-NTA agarose beads and microtubule pelleting was performed. Coomassie staining is presented. S stands for supernatant, P stands for pellet. (I) Effect of mNG-CCDC66 and mNG-CCDC66 (570-948) overexpression on microtubule organization and stability in cells. U2OS cells were transfected with mNG, GFP-CCDC66 or mNG-CCDC66 (570-948) expression plasmids. 24 h post-transfection, cells were treated with 5 µM nocodazole or 0.01% DMSO for 1 h, fixed with methanol and stained for the indicated proteins and DNA. Scale bar: 5 µm.

**Figure S6.**
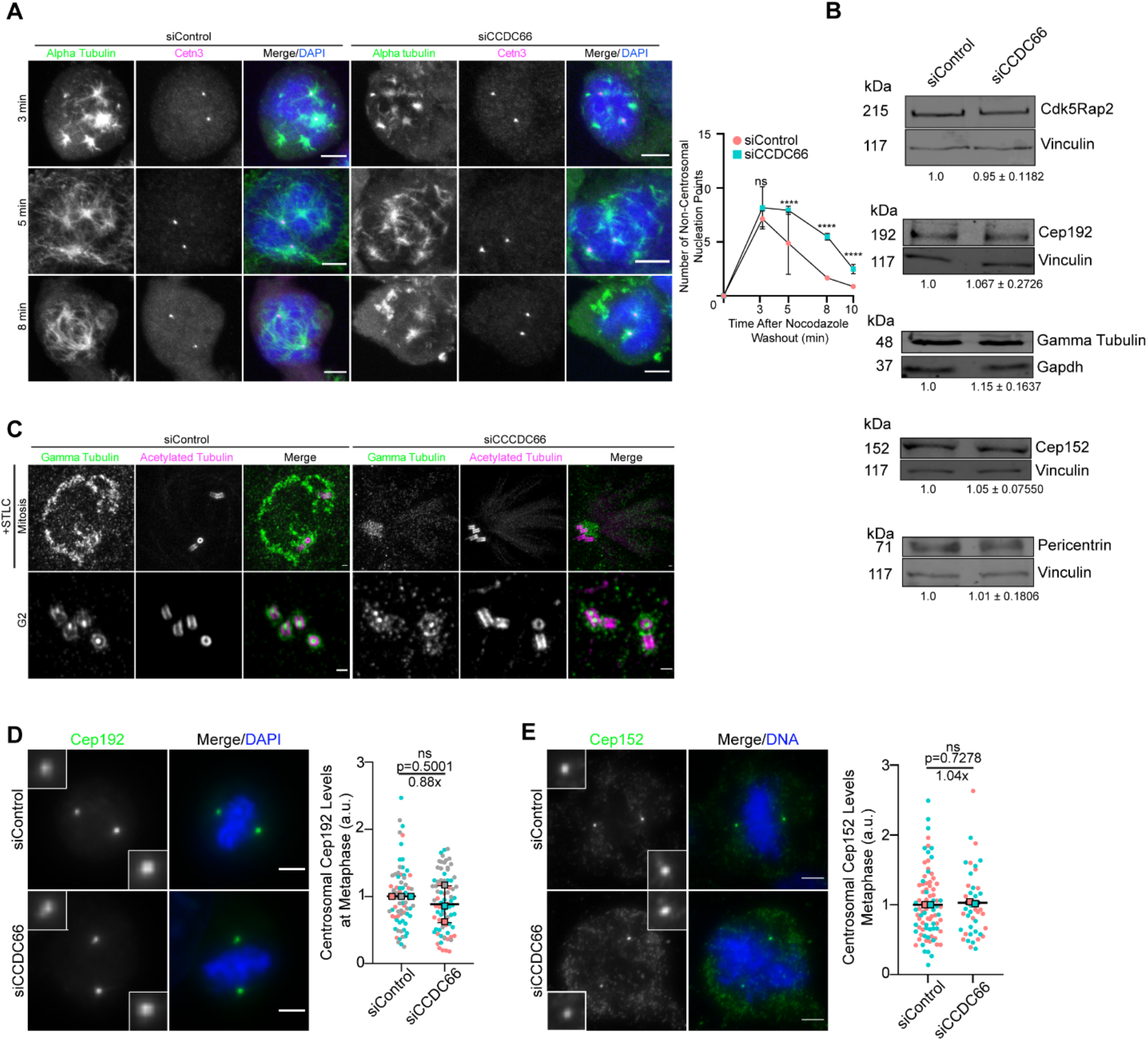
CCDC66 depletion comprimises centrosome maturation and microtubule nucleation. (A) Quantification of microtubule nucleation sites following nocodazole washout of STLC-synchronized control and CCDC66-depleted cells. Representative images are shown for control and CCDC66-depleted cells. The graph indicates the number of microtubule nucleation points that are counted from Centrin 3 and alpha-tubulin signals. Data represent the mean ±SEM of two independent experiments. (**p<0.01) Scale bar: 5 µm. (B) Effect of CCDC66 depletion on the cellular abundance of PCM proteins. U2OS cells were transfected with siControl or siCCDC66, and 48 h after transfection extracts from cells were immunoblotted for CDK5RAP2, CEP192, CeEP152, gamma-tubulin, Pericentrin and vinculin (loading control) or GAPDH (loading control). Band intensities were measured on ImageJ and normalized against background and vinculin intensities. Data represent the mean ±SEM of three independent experiments. (C) Ultrastructure expansion microscopy (U-ExM) analysis of control and CCDC66 depleted cells. U2OS cells were transfected with control and CCDC66 siRNA. 48 post-transfection, cells were synchronized by 16 h STLC treatment and prepared for imaging. Cells were stained for gamma-tubulin and acetylated tubulin, imaged using confocal microscopy and deconvolved. Mitotic and G2 cells were picked for representation. (D-E) Effects of CCDC66 depletion on abundance of PCM proteins at the spindle poles. U2OS cells were transfected with control and CCDC66 siRNA. After 48 h, cells were fixed with methanol and stained for (D) CEP192 and (E) CEP152. Centrosomal abundance of PCM proteins was measured as described in Fig. 4D. Images for each panel represent cells captured with the same camera settings from the same coverslip. Data represent mean ±SEM of two (CEP152) or three (CEP192) independent experiments. (ns: not significant). Scale bar: 5 µm

**Figure S7.**
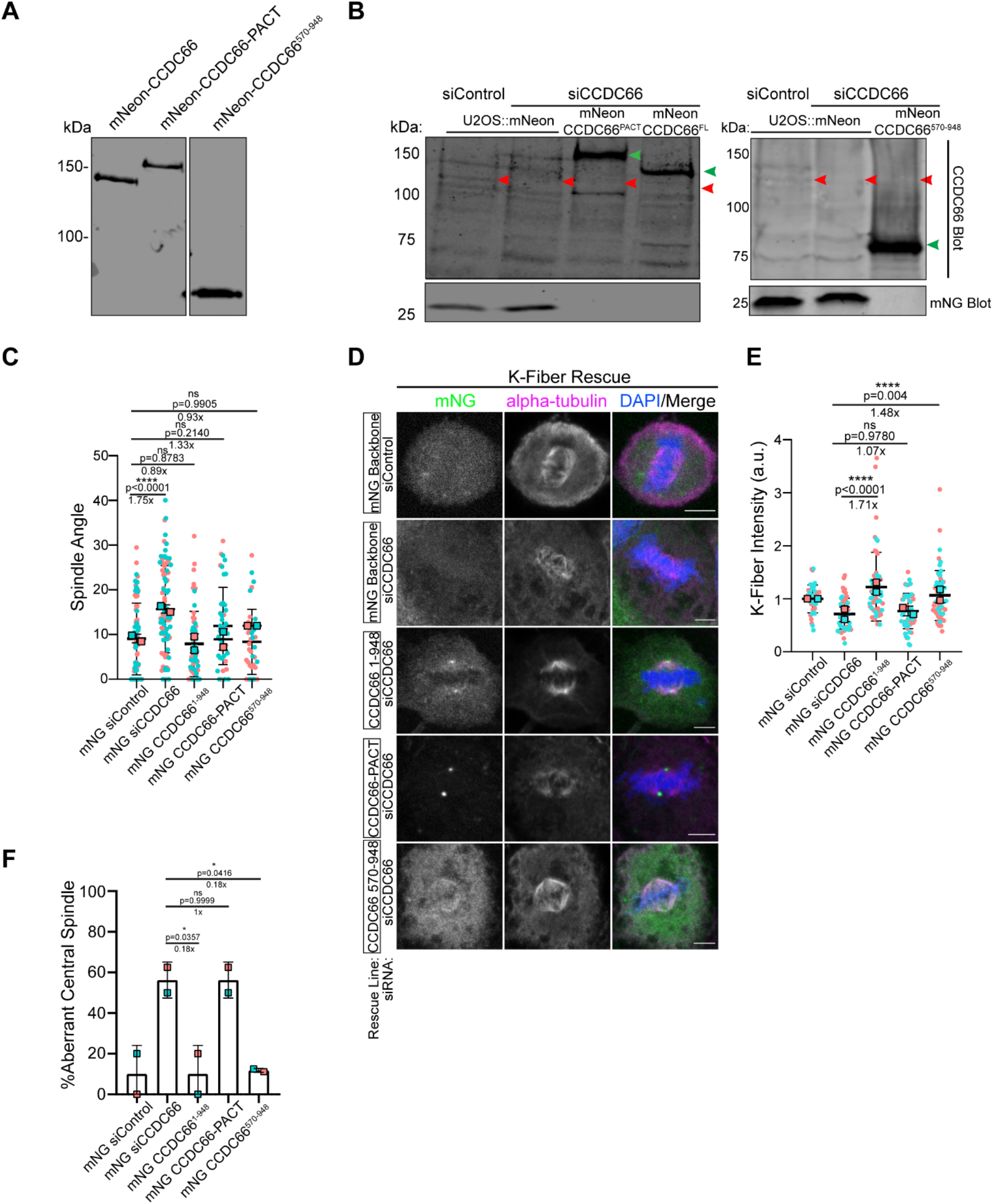
Phenotypic rescue for spindle positioning and k-fiber stability using stable lines expressing mNG-CCDC66 fusion constructs. (A) Validation of U2OS cells lines that stably express siRNA-resistant mNG-CCDC66, mNG-CCDC66-PACT and mNG-CCDC66 (570-948). Extracts from cells were prepared, resolved by SDS-PAGE and blotted with mNG antibody. (B) Validation of siRNA resistance of the CCDC66 rescue constructs. U2OS cells were transfected with control and CCDC66 siRNA. 48 h post-transfection extracts from cells were prepared, resolved by SDS-PAGE and blotted with CCDC66 antibody. The red arrowheads indicate endogenous CCDC66, which is masked due to higher expression of the fusion proteins and high background associated with the CCDC66 antibody. The green arrowheads indicate the mNG fusions of CCDC66. (C) Quantification of Figure 7A. Spindle angle was calculated by the formula α=180*tan^-1^(h/L)/π where h represents the stack difference between two spindle poles, L represents the distance between spindle poles. Data represent the mean ±SEM of two independent experiments. (**p<0.01 ***p<0.001 ****p<0.0001). (D) Representative images for the K-Fiber rescue experiment performed using U2OS::mNeonGreen, U2OS::mNeonGreen-CCDC66^1-948^, U2OS::mNeonGreen-CCDC66-PACT and U2OS::mNeonGreen-CCDC66^570-948^ stable cells. Cells were transfected with control and CCDC66 siRNA. 48 h post-transfection, they were fixed with methanol and stained for alpha-tubulin and DAPI. Scale bar: 5 µm. (E) Quantification of Figure 7D. Graph represents the intensity of K-fibers with mean ±SEM of two independent experiments. (****p<0.0001, ns: not significant). (F) Quantification of Figure 7D. Graph represents the percentage of aberrant central spindle with mean ±SEM of two independent experiments. (****p<0.0001, ns: not significant).

### Movie Legends

**Movie 1**. U2OS::mNG-CCDC66 cells were incubated with 100 nm SiR-Tubulin overnight and imaged with Leica SP8 Confocal microscopy equipped with an incubation chamber. Images are taken every 4 minutes. Scale bar: 5 µm.

**Movie 2**. mCherry-H2B::U2OS cells were transfected with non-targeting siRNA and imaged with Leica SP8 Confocal microscopy equipped with an incubation chamber with 20x objective. Images were taken every 6 minutes for 16 h. Scale bar: 5 µm.

**Movie 3**. mCherry-H2B::U2OS cells were transfected with CCDC66 siRNA and imaged with Leica SP8 Confocal microscopy equipped with an incubation chamber with 20x objective. Images were taken every 6 minutes for 16 h. This movie represents a CCDC66 depleted cell going through normal mitotic progression. Scale bar: 5 µm.

**Movie 4**. mCherry-H2B::U2OS cells were transfected with CCDC66 siRNA and imaged with Leica SP8 Confocal microscopy equipped with an incubation chamber with 20x objective. Images were taken every 6 minutes for 16 h. The cell marked with the asterisk represents a CCDC66 depleted cell going through apoptosis after delayed mitosis. Scale bar: 5 µm.

**Movie 5**. mCherry-H2B::U2OS cells were transfected with CCDC66 siRNA and imaged with Leica SP8 Confocal microscopy equipped with an incubation chamber with 20x objective. Images were taken every 6 minutes for 16 h. This movie represents a CCDC66 depleted cell going through premature mitotic exit. Scale bar: 5 µm.

**Movie 6**. mCherry-H2B::U2OS cells were transfected with CCDC66 siRNA and imaged with Leica SP8 Confocal microscopy equipped with an incubation chamber with 20x objective. Images were taken every 6 minutes for 16 h. This movie represents a CCDC66 depleted cell with prometaphase arrest and non-persistent metaphase plate. Scale bar: 5 µm.

**Movie 7**. mCherry-H2B::U2OS cells were transfected with CCDC66 siRNA and imaged with Leica SP8 Confocal microscopy equipped with an incubation chamber with 20x objective. Images were taken every 6 minutes for 16 h. This movie represents a CCDC66 depleted cell with a cytokinesis defect. Scale bar: 5 µm.

**Movie 8**. mCherry-H2B::U2OS cells were transfected with CCDC66 siRNA and imaged with Leica SP8 Confocal microscopy equipped with an incubation chamber with 20x objective. Images were taken every 6 minutes for 16 h. This movie represents a CCDC66 depleted cell with a cytokinesis defect. Scale bar: 5 µm.

## Supplemental Movie Legends

**Movie S1**. RPE1::mNG-CCDC66 cells were incubated with 100 nm SiR-Tubulin overnight and imaged with Leica SP8 Confocal microscopy equipped with an incubation chamber. Images are taken every 2 minutes. Scale bar: 5 µm.

